# Identification of distinct pathological signatures induced by patient-derived α-synuclein structures in non-human primates

**DOI:** 10.1101/825216

**Authors:** M. Bourdenx, A. Nioche, S. Dovero, M.-L. Arotcarena, S. Camus, G. Porras, M.-L. Thiolat, N. P. Rougier, A. Prigent, P. Aubert, S. Bohic, C. Sandt, F. Laferrière, E. Doudnikoff, N. Kruse, B. Mollenhauer, S. Novello, M. Morari, T. Leste-Lasserre, I. Trigo Damas, M. Goillandeau, C. Perier, C. Estrada, N. Garcia-Carrillo, A. Recasens, N. N. Vaikath, O. M. A. El-Agnaf, M. Trinidad Herrero, P. Derkinderen, M. Vila, J. A. Obeso, B. Dehay, E. Bezard

## Abstract

Dopaminergic neuronal cell death, associated with intracellular α-synuclein (α-syn)-rich protein aggregates (termed ‘Lewy bodies’), is a well-established characteristic of Parkinson’s disease. Much evidence, accumulated from multiple experimental models has suggested that α-syn plays a role in PD pathogenesis, not only as a trigger of pathology but also as a mediator of disease progression through pathological spreading. Here we have used a machine learning-based approach to identify unique signatures of neurodegeneration in monkeys induced by distinct α-syn pathogenic structures derived from PD patients. Unexpectedly, our results show that, in non-human primates, a small amount of singular α-syn aggregates is as toxic as larger amyloid fibrils present in the LBs, thus reinforcing the need for preclinical research in this species. Furthermore, our results provide evidence supporting the true multifactorial nature of PD as multiple causes can induce similar outcome regarding dopaminergic neurodegeneration.

## Introduction

The seminal work of Braak and colleagues suggesting that Lewy body (LB) pathology follows a predictable pattern of progression within the brain in Parkinson’s disease (PD) (*1*) as well as the ‘host-to-graft’ observation (*2*–*4*) led to the development of experimental models based on injection with α-synuclein (α-syn – the primary protein component of LB) assemblies (*5*–*7*). These experimental models suggest that α-syn, in pathological conformations such as the one found in LBs, initiates a cascade of events leading to dopaminergic neuron degeneration as well as cell-to-cell propagation of α-syn pathology through a self-templating mechanism.

Several studies have suggested that pre-fibrillar oligomers may represent one of the major neurotoxic entities in PD (*8*, *9*). This notion has been derived primarily from studies using large doses of recombinant α-syn applied to cell cultures or injected into adult mice, over-expressing either mutant or wild-type α-syn (*10*). In agreement with these findings, we have shown that intracerebral injection of low doses of α-syn-containing LB extracts, purified from the substantia nigra, pars compacta (SNpc) of postmortem PD brains, promotes α-syn pathology and dopaminergic neurodegeneration in wild-type mice and non-human primates (*11*). Importantly, this neuropathological effect was directly linked to the presence of α-syn in LB extracts, since immuno-depletion of α-syn from the LB fractions prevented the development of pathology following injection into wild-type mice.

In this study, our aim was to thoroughly investigate this experimental model of synucleinopathy in non-human primates. The initial study design was to administrate fractions derived from the same PD patients containing either soluble and small α-syn aggregates (hereafter named noLB) or LB-type aggregates (hereafter named LB). However, because of the unexpected finding that non-human primates, unlike mice, are susceptible to soluble or finely granular α-syn, we sought to elucidate the response characteristics induced by either LB or noLB fractions. To achieve a thorough analysis of these α-syn-related characteristics, we took advantage of the strength of machine-learning algorithms for discovering fine patterns among complex sets of data and developed a new method compatible with the constraints of experimental biology. We here report the identification of primate-specific responses to selected α-syn assemblies associated with different pathogenic mechanisms. Overall, our results support the concept of the multifactorial nature of synucleinopathies.

## Results

### Purification and characterization of α-synuclein extracts from PD patients

NoLB and LB fractions were obtained from the SNpc of five sporadic PD brains exhibiting conspicuous LB pathology. The samples were processed through differential ultracentrifugation in a sucrose gradient, and analyzed for the presence of α-syn aggregates by filter retardation assay (Fig. 1A) (*11*). Further characterization of noLB and LB fractions was performed by co-localization of α-syn and the amyloid dye Thioflavin S (Fig. 1B) as well as ultrastructural examination by electron microscopy (Fig. 1C). These assays confirmed the presence of misfolded α-syn in both fractions. We also performed biochemical characterization of the stability of assemblies after proteinase K digestion (Fig. 1D) and detergent treatments (Fig. 1E) followed by α-syn dot-blot assays. While total α-syn content was comparable between selected fractions (as measured by α-syn ELISA), LB fractions showed higher resistance to proteinase K treatment (noLB t_1/2_=15.23minutes vs LB t_1/2_>60minutes) (Fig. 1D) as well as greater resistance to multiple detergents, including 8M Urea (Fig. 1E). We then measured the content of α-syn aggregates using human α-syn aggregation TR-FRET-based immunoassay, which revealed a significantly higher amount of aggregated α-syn in LB fractions (Fig. 1F). To obtain insight into the content of monomeric and aggregated α-syn within noLB and LB fractions of PD patients, sarkosyl treatment was applied to both fractions to induce physical separation, and then velocity sedimentation and density floatation gradients were performed to quantify these two respective populations and determine their relative abundance in each fraction (Fig. S1 A-H). Strikingly, while LB fractions contained ~90% of aggregated α-syn, noLB fractions were composed of ~10% of this pathological form of the protein (Fig. S1 I). Also, in order to confirm the quality of the LB extraction, we performed a filter retardation assay which showed that LB fractions, but not noLB fractions, were highly enriched in known components of LBs, such as phosphorylated S129 α-syn, ubiquitin, p62, hyperphosphorylated tau and Aβ (Fig. S2 A).

**Fig. 1.**
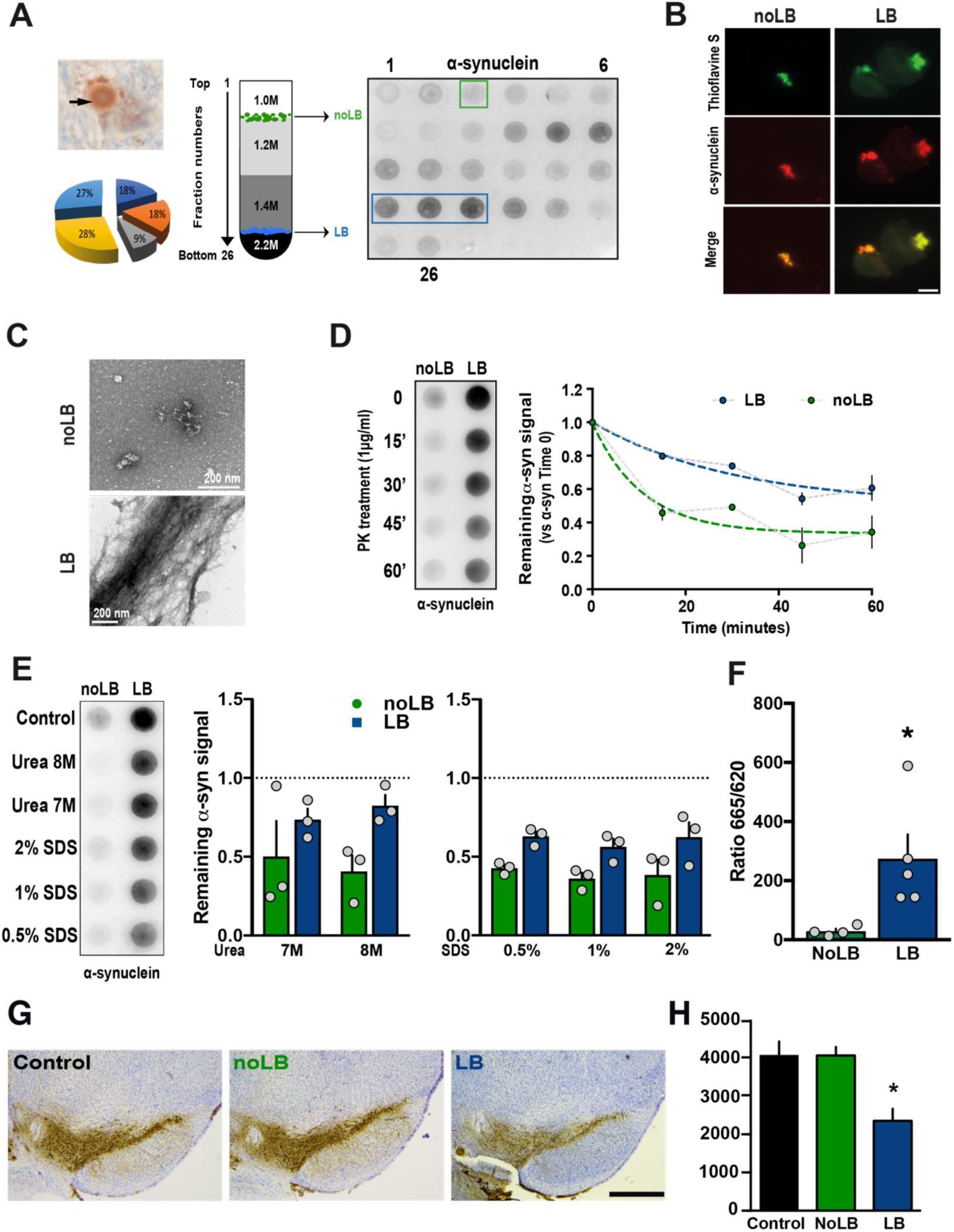
Purification and characterization of Lewy bodies (LB) and noLB inocula from Parkinson disease (PD) brains. (**A, left**) Immunohistochemistry image of α-synuclein–positive LB (arrows) in nigral postmortem brain samples (PD #1; α-synuclein in brown, neuromelanin in dark-brown) before sucrose gradient purification. The pie chart indicates the relative contribution of the 5 patients to the final pool of LB and noLB inocula (**A, middle**) Schematic representation of the sucrose gradient fractionation procedure used to purify LB/noLB-containing fractions from freshly frozen postmortem nigral brain tissue of 5 sporadic PD patients. (**A, right**) Filter retardation assay probed with a human α-synuclein antibody to assess the presence of α-synuclein aggregates in the different fractions obtained by sucrose gradient fractionation from freshly frozen postmortem nigral brain tissue from sporadic PD patients (PD #1). Green rectangle indicates noLB-containing fraction and blue rectangle highlights LB-containing fraction selected to prepare the mixture used for injections. (**B**) Confocal examination of purified noLB and LB fractions with α-syn immunofluorescence (red) and thioflavin S staining (green). Both LB and noLB present thioflavin S-positive aggregates but much smaller in noLB fractions. Scale bar = 10μm. (**C**) Ultrastructural examination of noLB and LB fractions by electron microscopy showing massive fibrils in LB fractions while noLB fractions contain, besides soluble α-syn, some punctiform small size aggregates. (**D**) NoLB and LB fractions derived from PD brains (left panel) were treated with 1 μg/ml proteinase K for 0, 15, 30, 45 and 60 min and analyzed by immunoblotting with syn211 antibody. The EC50 value was determined as the concentration at which this ratio is decreased by 50%. The corresponding EC50 value for LB (>60 min) was approximately fourfold greater than with noLB (15.23 min) **(E**) NoLB and LB fractions were treated for 6h with increasing concentrations of either urea or SDS or buffer as control. Syn211 was used to detect the forms of α-synuclein. The LB fractions appear to be more resistant to breakdown compared with noLB fractions in both urea (F_(1,8)_=6.063, p=0.0392) and SDS treatments (F_(1,12)_=17.41, p=0.0013). The dotted line show levels of control fractions. Comparison were made using Two-Way ANOVA. **(F**) TR-FRET immunoassay analysis of noLB and LB fractions. Fluorescence measurements were taken 20h after antibody. Analysis by unpaired Student’s t-test (t_(7)_=2,623, p=0,0343). *: P<0.05. Mean ± SEM, n=4-5. (**G**) Representative pictures of tyrosine hydroxylase (TH)-positive substantia nigra pars compacta (SNpc) neurons (brown; Nissl staining in purple) in non-injected, noLB or LB-injected mice at 4 months after injections. Scale bars=500μm. (**H)** Quantification of TH-positive Substantia Nigra pars compacta (SNpc) neurons by stereology in control, LB- and noLB-injected mice. Control mice, n=10, LB-injected mice at 4 months, n=10, No-LB-injected mice at 4 months, n=10. One-way ANOVA followed by Tukey test for multiple comparisons. *: p<0.05 compared with control and noLB-injected side at 4 months.

Micro-Infrared Spectroscopy of LB and noLB fractions was performed to show conformational changes in amyloid structures at the molecular level (Fig. S2 B-E) and this confirmed the presence of α-sheet structures in both assemblies (Fig. S2 B-C). Although their velocity of sedimentation and density floatation characteristics were similar, the aggregates present in the LB and noLB fractions were different in nature based upon the evidence of Micro-Infrared Spectroscopy. Principal component analysis (PCA) showed that, in the LB fractions, large aggregates corresponding to the major pieces of LB were present (Fig. S2D, cluster on the right). PCA further showed that, in the range of 1,590-1,700 cm^−1^, the LB group contained a fraction of amyloid aggregates with different amyloid structures from those in the noLB group as they clearly segregated by PCA in two clusters (Fig. S2 D-E). Altogether, these results suggest that while LB fractions primarily contained large aggregated α-syn fibrils, noLB fractions contained soluble α-syn and a smaller enrichment of α-syn aggregates featuring a specific amyloid structure not found in the LB fractions.

Data from several studies suggest that both recombinant α-syn preformed fibrils (*12*–*14*) and patient-derived α-syn (*11*) can promote pathogenic templating of endogenous α-syn ultimately leading to dopaminergic neurodegeneration in SNpc. Following quantification by ELISA, both mixes of fraction were diluted to ~24 pg α-syn per microliter. Then, those fractions were tested for their pathogenic effects on TH-positive dopaminergic neurons in primary mesencephalic cultures (Fig. S3 A) as well as *in vivo* in wild-type mice. Four months after supranigral injection, LB-injected mice displayed, as expected, significant dopaminergic degeneration, while noLB injections in mice had no impact on dopaminergic neurons (Fig. 1G-H) as we have previously reported for other SNpc-derived LB fractions (*11*), thus validating the toxicity of the preparation prior to injection into non-human primates.

### Intrastriatal injection of LB and noLB fractions from Parkinson’s disease patients induces nigrostriatal neurodegeneration in baboon monkeys

To determine the mechanisms of α-syn aggregates toxicity in a species closer to humans, adult baboon monkeys (n=4-7 per experimental group) received bilateral stereotaxic injections (100μl) of either LB or noLB fractions into the putamen before euthanasia 24 months post-injection. This time-frame was chosen based on our previous studies indicating that after 14 months post-injection, ongoing pathogenic effects can already be measured, and was extended to potentially reach disease-relevant lesions. Two years after administration, LB-injected monkeys displayed significant striatal dopaminergic terminal loss both in the putamen and in the caudate nucleus, accompanied by a significant decrease in tyrosine hydroxylase (TH) immunoreactivity in the substantia nigra pars compacta (SNpc) (Fig. 2). Stereological counts showed that LB-injected animals exhibited TH-positive and Nissl-positive cell loss in the SNpc (16% and 23%, respectively). No overt parkinsonism was observed, however, since the extent of the lesion remained below the threshold for symptom appearance; i.e. 45% of cell loss (*15*), compared to an age-matched control group.

**Fig. 2.**
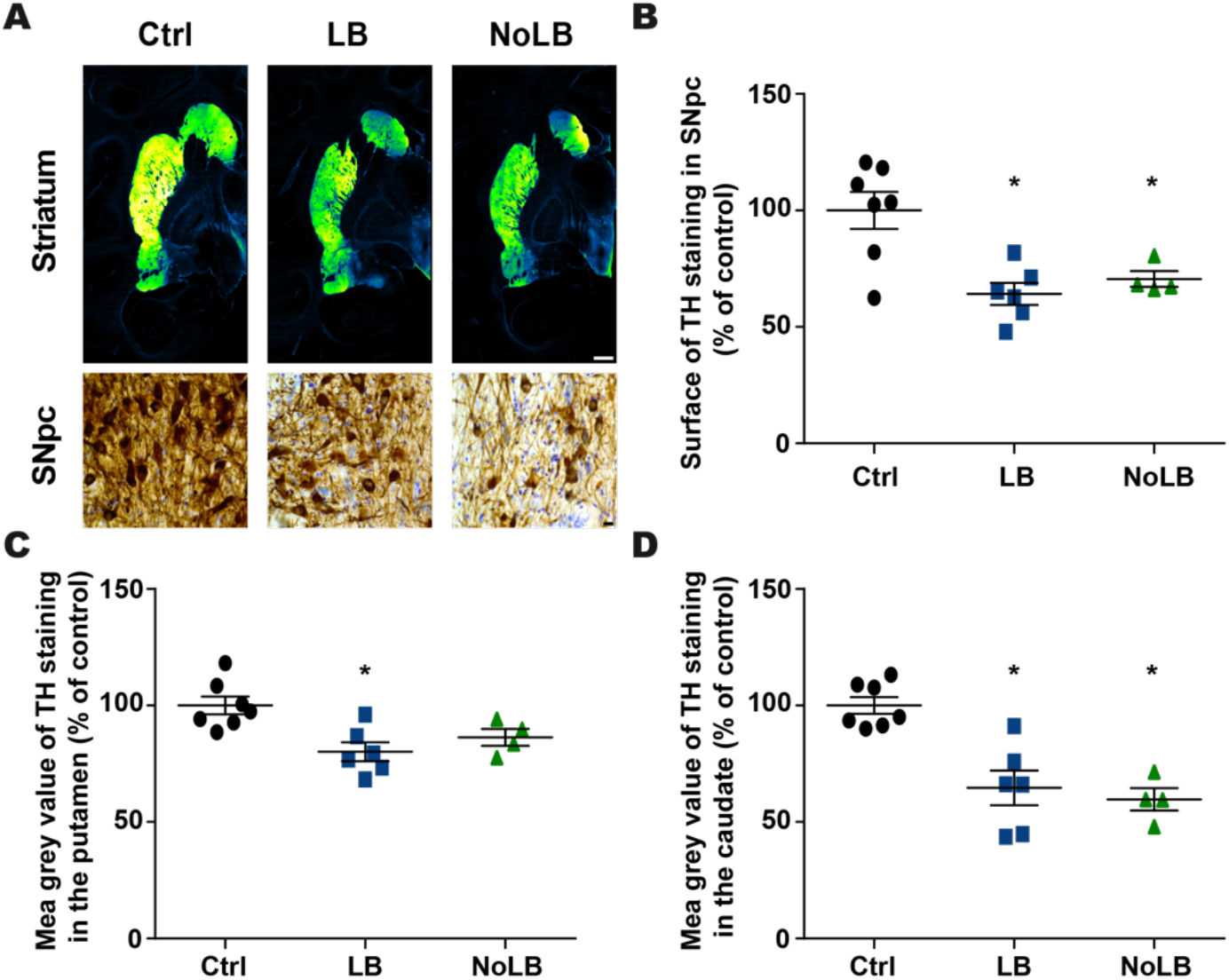
Intrastriatal injection of Lewy bodies (LB) and noLB fractions from Parkinson’s disease patients induces nigrostriatal neurodegeneration in baboon monkeys. (**A**) Tyrosine hydroxylase (TH) staining at striatum and Substantia Nigra pars compacta (SNpc) levels. A green fire blue LUT (lookup table) was used to enhance contrast and highlight the difference between non- injected, LB-injected and noLB-injected baboon monkeys at striatum level. Scale bars = 5mm (striatum) and 10μm (SNpc). (**B**) Scatter plot of TH immunostaining in SNpc. F_(2,14)_=9.439, p=0.0025. Control vs LB-injected: p=0.0029. Control vs noLB-injected: p=0.0248. (**C, D**) Scatter plots of mean grey values of striatal TH immunoreactivity in the putamen (F_(2,14)_=7.313, p=0.0067; Control vs LB-injected: p=0.0059) (**C**) and in the caudate (F_(2,14)_=16.25, p=0.0002; Control vs LB-injected: p=0.0008; Control vs noLB-injected: p=0.0008) (**D**) in non-injected, LB-injected and noLB-injected baboon monkeys. The horizontal line indicates the average value per group ± SEM (n=7 from control animals; n=6 for LB-injected animals; n=4 for noLB-injected animals). Comparison were made using One-Way ANOVA and Tukey’s correction for multiple comparison. *p< 0.05 compared with control animals.

At odds with mice either generated for the purpose of this study (Fig. 1G-H), previously published (*11*), or produced in the context of other in-house studies (data not shown), noLB-injected monkeys showed degeneration of the nigrostriatal pathway including dopaminergic cell loss (i.e. 16% of TH-positive neurons and 28% of Nissl-positive neurons quantified by stereology), similar to that observed in LB-injected monkeys (Fig. 2). Facing such an unexpected finding, we aimed to identify specific characteristics of the pathological mechanisms involved in α-syn toxicity induced by each fraction independently, using a large-scale approach in combination with machine learning for pattern identification.

### Machine-learning algorithm predicts nigrostriatal degeneration

We performed an exploratory approach and aimed to distinguish relevant variables allowing accurate prediction of neurodegeneration (i.e., to operate a feature selection). Overall, we investigated a large number of variables tapping on behavioral, histological, biochemical, transcriptional and biophysical approaches (Fig. 3A) applied to several brain areas (n=40 – Fig. 3B), totalizing 180 variables measured for each individual (Fig. S4A for variable abbreviation nomenclature; Table S1 for exhaustive list of variables; Table S2 features all raw data). We first extracted from this dataset, every variable that actually quantified neurodegeneration (i.e. dopaminergic markers such as TH or dopamine transporter by immunohistochemistry), ending up with 163 variables per animal.

**Fig. 3.**
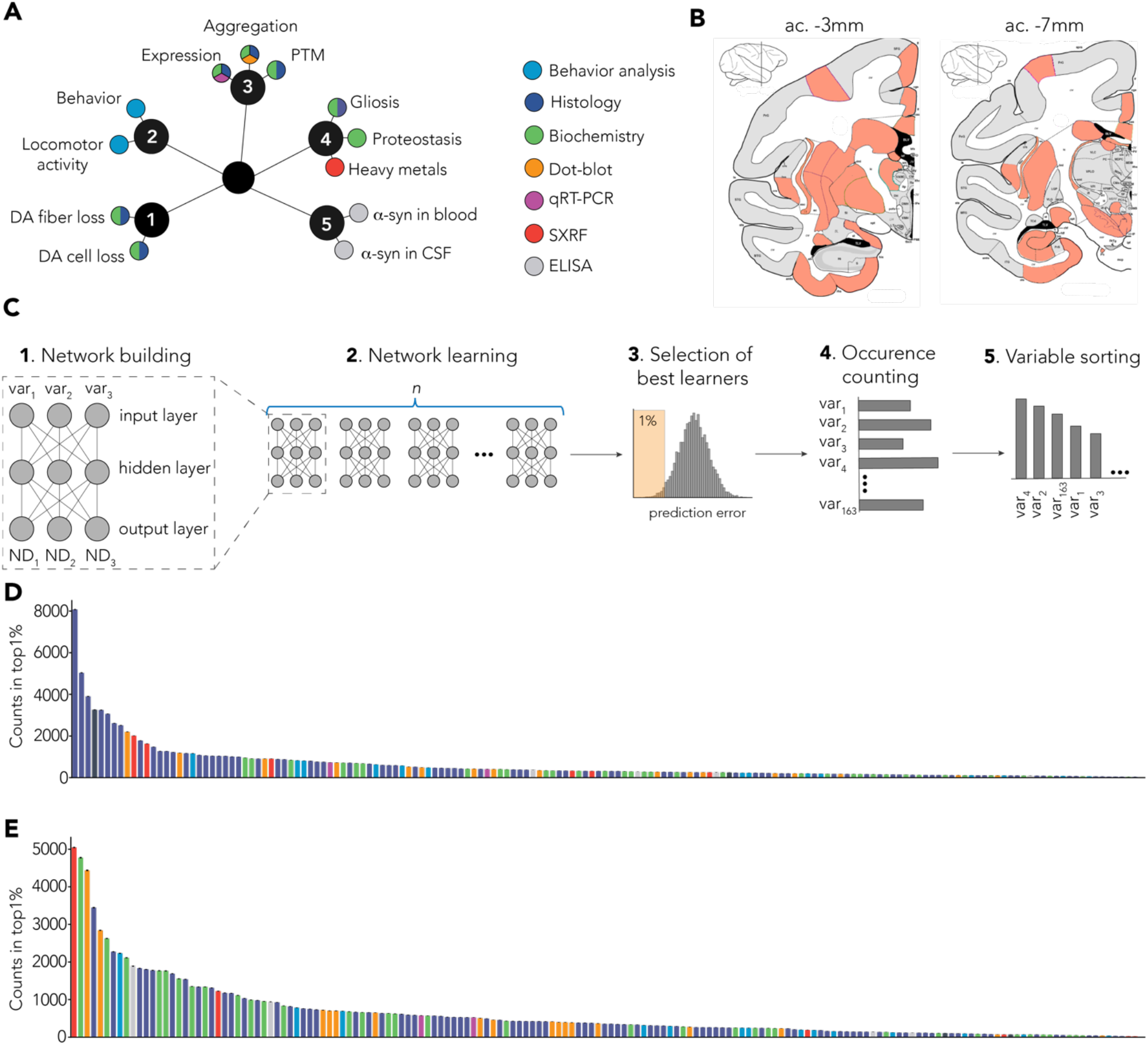
Multiple-layer perceptron (MLP)-based identification of specific signature. (**A**) Several endpoints (n=180) were measured using multiple methods (colors). Endpoints can be grouped as clusters: *1*. Dopaminergic degeneration, *2*. Behavior, *3*. α-syn-related pathology. *4*. Non-α-syn related pathology. *5*. Putative biomarkers. (**B**) Multiple brain regions (n=40) were investigated from coronal sections at 2 levels: anterior commissure (ac.) −3mm (striatum, entorhinal cortex) and −7mm (SNpc, hippocampus). (**C**) Detailed methodology. *1*. Representative scheme of one MLP predicting 3 neurodegeneration-related variables (ND_1_, ND_2_, ND_3_) with 3 experimental variables as input (var_1_, var_2_, var_3_). Out of the 180 variables measured in total, 163 were used as inputs for the MLP. *2*. One MLP was trained for every unique combination of 3 variables. *3*. Combinations were ranked based on their prediction error and top1% were selected for further analysis. *4*. Combinations were deconvoluted to extract single variables and count occurrence of individual variables. *5*. Variables were sorted based on the number of occurrences in the top1% of the best combination. (**D**) Raw ranking obtained for LB-injected animals. Color code highlights measurement methods as in A. (**E**) Raw ranking obtained for noLB-injected animals. Color code highlights measurement methods as in A.

Then, to operate feature selection, we designed a distributed algorithm using multiple layer perceptron (MLP) (Bourdenx and Nioche, 2018), a classic machine-learning algorithm based on artificial neural network that is able to approximate virtually any functions (Hornik et al., 1989). This algorithm was given, as input, the data obtained for each animal for the 163 aforementioned variables and its output is a rank of these variables regarding their ability to predict three indicators of dopaminergic tract integrity; that were levels of tyrosine hydroxylase staining in (i) the SNpc, (ii) the putamen and (iii) the caudate nucleus.

The main difficulty was to overcome the large number of input variables (163) compared to the sample size (n=4-7 per group), which can induce a selection and reporting bias (Kuncheva and Rodriguez, 2018). In order to tackle this “p > n” problem, instead of using a single network that could be prone to overfitting, we put in competition several networks.

Each MLP was composed of a single hidden layer of 3 neurons (Fig. 3C). It has as input a subset of 3 variables (out of the 163) and as output the 3 indicators of dopaminergic tract integrity. In total, we used 708,561 sets of 3 inputs variables. Every instance of MLP was trained with 80% of our sample (always a combination of control and injected animals) and tested on the remaining 20%. The performance of each set of 3 input variables was evaluated according to the difference between the predicted values of TH staining and the actual ones.

We focused on the top 1% of the best networks and counted the occurrence of each of the 163 variables in the subset of 3 variables used by these best networks (Fig. 3C). We ranked each variable according to the number of occurrences (Fig. 3C) for LB- (Fig. 3D) and noLB-injected animals (Fig. 3E) independently.

In order to avoid possible overfitting, we used several methods in combination. First, we performed cross-validation by splitting the dataset into two parts: a training and a testing set of data. 80% of the data were randomly selected to train the networks (and independently for each network), while the 20% remaining were used to evaluate the networks. Then, in order to evaluate the robustness of the quality of prediction for a given set, we repeated this cross-validation step 50 times for every set of 3 input variables (each network was trained and tested using a different partition of the dataset - total number of network: 35,428,050). Lastly, we generated random data and used them as input for the MLP. As expected, performances were significantly lower compared to our actual dataset (Fig. S4B, C).

Overall, this unique approach allowed us to rank input variables according to their explanatory power and therefore to extract the strongest predictors of neurodegeneration for each experimental group. Interestingly, despite similar levels of nigrostriatal degeneration between LB- and noLB-injected animals (Fig. 2B), the algorithm allowed us to identify differential variable sorting patterns (Fig. 3D-E).

### MLP-derived signatures can identify unique characteristics between experiment group

Next, we compared the LB and noLB characteristics using the rank-rank hypergeometric overlap (RRHO) test (Fig. 4A). Interestingly, low similarity was observed for the highly ranked variables suggesting specific differences in the biological response to the injection of LB or noLB (Fig. 4B). Focusing on the 20 first variables that showed low similarity between groups, we found that LB-exposed monkeys were characterized by both quantitative and qualitative changes in α-syn levels (i.e. phosphorylation at Ser129 and aggregation) especially in cortical areas corroborated by distinct methodologies as well as by a dysfunctional equilibrium in neurochemistry of basal ganglia output structures classically associated with parkinsonism (*16*, *17*) (Fig. 4C – Fig. S5). Conversely, noLB-exposed monkeys exhibited more diverse nigrostriatal-centric characteristics with variables related to α-syn aggregation, proteostasis and Zn homeostasis (Fig. 4D - Fig. S6). Together, we identified specific properties for both groups with limited overlap (35% −7/20 variables) for an identical level of degeneration.

**Fig. 4.**
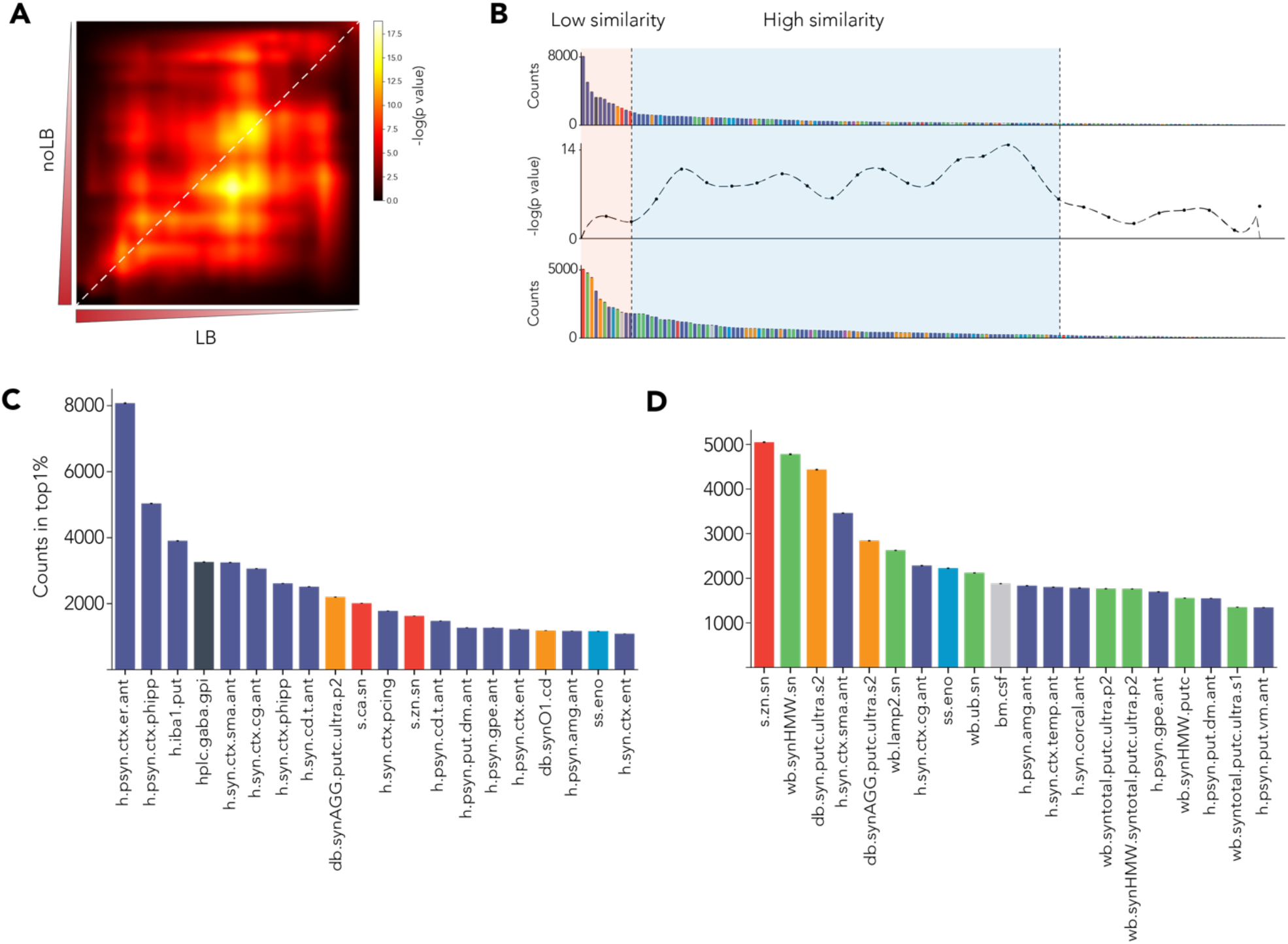
Direct comparison of MLP-derived signatures shows specific pattern between experiment groups. (**A**) Rank-rank hypergeometric overlap (RRHO) test between variable sorting of LB and noLB-injected animals. Highly enriched variables are in the lower left corner. Diagonal (highlighted by a red dashed line) was extracted to do a bin-to-bin comparison between LB and noLB signatures. (**B**) Signatures were aligned with RRHO and show low similarity in highly enriched variables (light orange background) and higher similarity for lower rank variables (pale blue background). (**C, D**) First 20 enriched variables for both LB-injected animals (**C**) and noLB-injected animals (**D**). Color code is similar to Fig. 2A. Detailed of variable names can be found in **Table S1**. Bars are mean +/−99% confidence interval estimated by bootstrap.

### Retrospective literature search validates MLP derived signatures

We next used a retrospective analysis to validates the relevance of the MLP-derived signature in PD. Although, some variables have never been investigated in the context of PD, others have been studied and reports exists in the literature. For instance, the amount of phosphorylated Ser129 α-synuclein in the entorhinal (*h.psyn.ctx.er.ant*) and parahippocampal (*h.psyn.ctx.phipp)* cortex - 1^st^ and 2^nd^ best predictors for the LB group – have been already associated with PD pathology. Studies of post-mortem brains from PD patients revealed the presence of LB in these regions which was correlated with disease progression(*18*) and predicted cognitive deficit in PD patients (*19*). Interestingly, the anterior entorhinal cortex has also been shown to be affected by severe α-syn pathology, related to olfactory dysfunction in prodromal phases of PD pathology (*20*). In addition, increased of levels of phosphorylated Ser129 α-syn in sensorimotor (*h.syn.ctx.sma.ant*) and cingulate cortices (*h.syn.ctx.cg.ant*), shared by both LB and noLB signatures, have already been reported by our group in an independent cohort of non-human primates (*11*).

Both LB and noLB signatures, and especially noLB, showed that variables related to α-syn aggregation status were among the best predictors (LB: 1 in top10 best predictors; noLB 3 in top10 best predictors). This was highly expected from the literature as α-syn aggregation has been associated with PD pathology (*21*).

Variables related to the proteostasis network (levels of the lysosomal receptor LAMP2 – *wb.lamp2.sn* - 6^th^ or amount of ubiquinated proteins – *wb.ub.sn* – 9^th^) were more specifically associated with the noLB signature. This is of high interest as proteostasis defect is more and more considered as a key step in pathogenicity (*22*–*24*).

Levels of the microglia marker, IbaI, was ranked as the third best predictor of neurodegeneration in the LB signature. Microglial inflammatory response was shown to be implicated in neurodegeneration in many animal models, including α-syn overexpressing and toxin-based animal model of PD (*25*).

Lastly, *postmortem* analysis of Zn^2+^ concentration in the brains of PD patients has shown elevated levels in the striatum and SNpc (*26*). Conversely, a recent meta-analysis showed a decrease of circulating Zn^2+^ levels in PD patients (*27*). In experimental models of PD, Zn^2+^ accumulation has been associated with dopaminergic degeneration in rodent exposed to mitochondrial toxins (*28*, *29*).

### Experimental confirmation of MLPs’ prediction

We aimed to confirm the relevance of the top first MLP selected variables. Since the LB signature was associated with changes in α-syn phosphorylation in cortical areas, we analyzed side-by-side the levels of α-syn and phosphorylated Ser129 α-syn in 18 brain regions (Fig. 5A). Interestingly, in agreement with the LB signature obtained from the MLP, LB-injected monkeys displayed a stronger accumulation of phosphorylated Ser129 α-syn compared to noLB-injected animals (Fig. 5A-B). Also, the 2 most enriched variables of the LB signature (i.e. phosphorylated α-syn levels in parahippocampal and entorhinal cortices (Fig. 4C)) showed significant negative correlations with degrees of degeneration (Fig. 5C-D), thus confirming their ability to predict neurodegeneration.

**Fig. 5.**
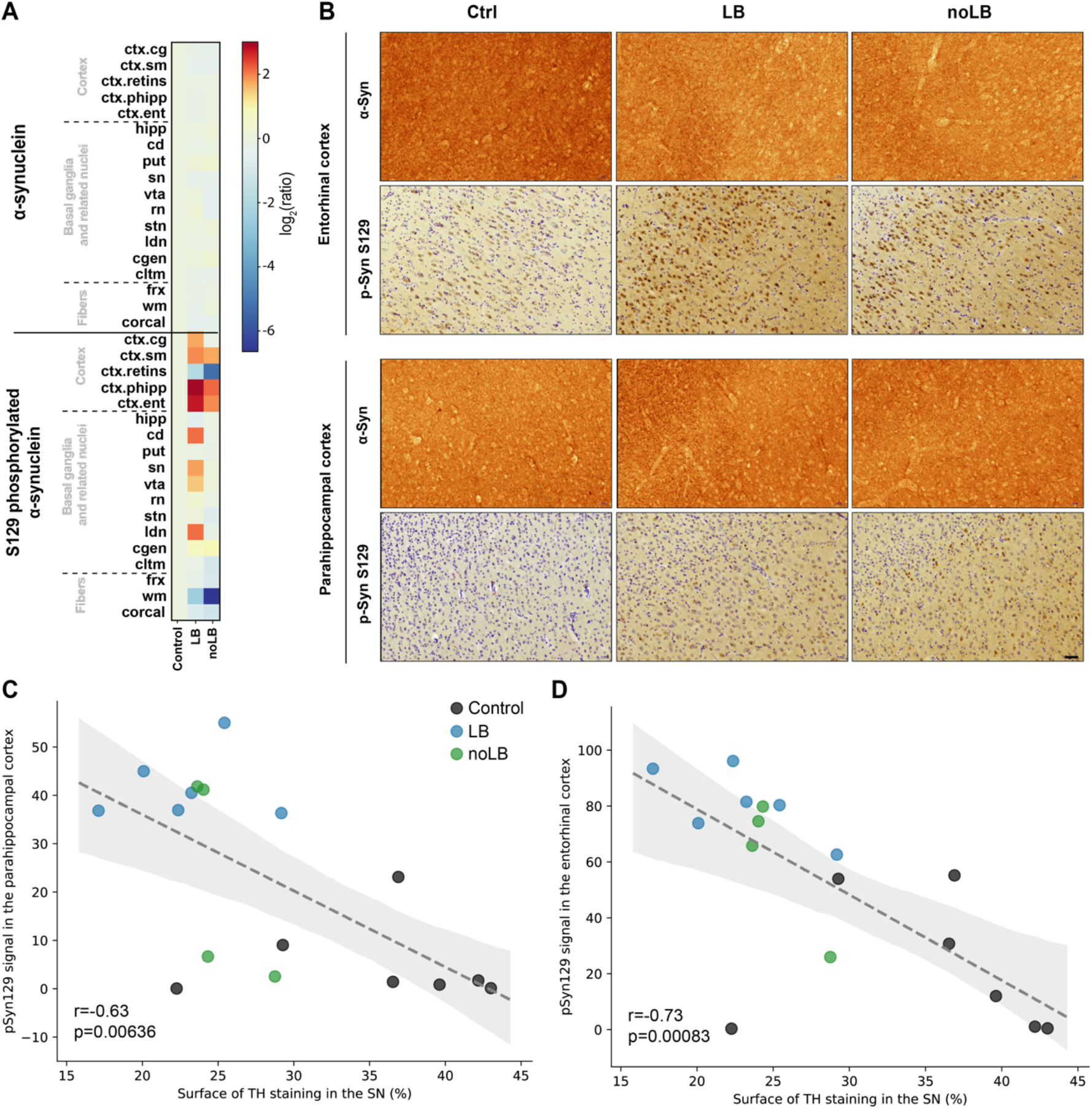
Levels of α-synuclein and phosphorylated α-synuclein in different brain regions. (**A**) Heat map representing the surface of α-synuclein (α-syn) and S129 phosphorylated α-syn immunostaining intensity in the brain of non-inoculated, LB-inoculated and noLB-inoculated baboon monkeys. The heat maps show all brain regions measured and are organized according in 3 main groups: cortical, basal ganglia and sub-cortical areas. From top to bottom: cingulate cortex (*ctx.cg*), sensorimotor cortex (*ctx.sm*), retro-insular cortex (*ctx.retins*), parahippocampal cortex (*ctx.phipp*), entorhinal cortex (*ctx.ent*), hippocampus (*hipp*), caudate nucleus (*cd*), putamen (*put*), substantia nigra (*sn*), ventral tegmental area (*vta*), red nucleus (*rn*), subthalamic nucleus (*stn*), lateral dorsal nucleus (*ldn*), lateral geniculate nucleus (*cgen*), claustrum (*cltm*), fornix (*frx*), white matter (*wm*), corpus callosum (*corcal*). The color bars represent the log2 value of the ratio of each brain regions. (**B**) Representative pictures of α-syn α-syn) and phosphorylated α-syn (pSyn S129) staining in the entorhinal and parahippocampal cortices. (**C, D**) Correlation between levels of phosphorylated α-syn in the parahippocampal cortex (**C**) and the entorhinal cortex (**D**) with levels of TH staining in the substantia nigra. Dotted line indicates the linear regression. Gray area indicates the 95% confidence interval around of the linear regression.

Then, we decided to confirm the relevance of one of the strongest predictors, the levels of Zn^2+^ in the SNpc in independent experiments. First, we observed a significant increase of Zn^2+^ in noLB-injected mice compared to sham-injected or LB-injected mice (Fig. S7A). Second, we analyzed the levels of Zn^2+^ in LB-injected macaque monkeys from a previous study of our laboratory (*11*). Interestingly, despite the fact that these experiments were done in a different non-human primate sub specie, injection of LB in the putamen (similar to the present study) or above the SNpc (different from the present study) induced elevation of Zn^2+^ levels in the SNpc, as measured by SR-XRF (Fig. S7B). Of note, the dimension of the effect was similar across studies (Fig. S7E). Then, to understand whether that modulation Zn^2+^ levels was specific to our experimental paradigm, we measured Zn^2+^ levels in the context of adeno-associated virus-mediated overexpression of mutant human α-syn in both rats and marmoset monkeys (*30*) using the same methodology (Fig. S7C, D). Here, overexpression of α-syn did not triggered accumulation of Zn^2+^ in the SNpc (despite inducing dopaminergic neurodegeneration – (*30*) suggesting that this phenomenon is specific to seeding experiment paradigms.

Lastly, we analyzed a publicly available cortical proteomic database of healthy individual and PD patients. Of interest, we observed that several Zn^2+^ transporters were elevated in the brains of PD patients thus suggesting a zinc dyshomeostasis in patients (Fig. S7F). Indeed, plasma membrane transporters such as the zinc transporter 1 (ZnT1), the Zrt-/Irt-like protein 6 (ZIP6) and ZIP10 showed increased levels (Fig. S7G-I) while the synaptic vesicle membrane transporter ZnT3 remained constant (Fig. S7J).

### Association metric shows independence of strong predictors

As we used combinations of 3 variables and because of the structure of MLPs, one could expect that some combinations would complement each other to allow finer prediction of neurodegeneration levels. To address this question, we used a classic measurement of association in the field of data-mining: lift (*31*) and plotted the results as network plots showing association (edge size) and enrichment in the best learners (node size). Lift calculation was corrected for error prediction to avoid detrimental association between variables. The first observation was that the most enriched variables (top 3 to 5) appeared to be self-sufficient to predict the neurodegeneration levels with minimal error (Fig. 6). Some variables, with modest enrichment, showed strong positive associations that were specific to each experimental group. Associated variables in LB-injected monkeys were: (i) α-syn-related parameters along the SNpc-striatum-cortex axis, an impairment of locomotion and the ethologically-defined orientation of the animals towards their environment (Fig. 6 top left inset); (ii) oligomeric α-syn species measured in the midbrain and striatum equally associated, but to lesser extent, with α-syn levels in cortex and plasma (Fig. 6 top right inset).

**Fig. 6.**
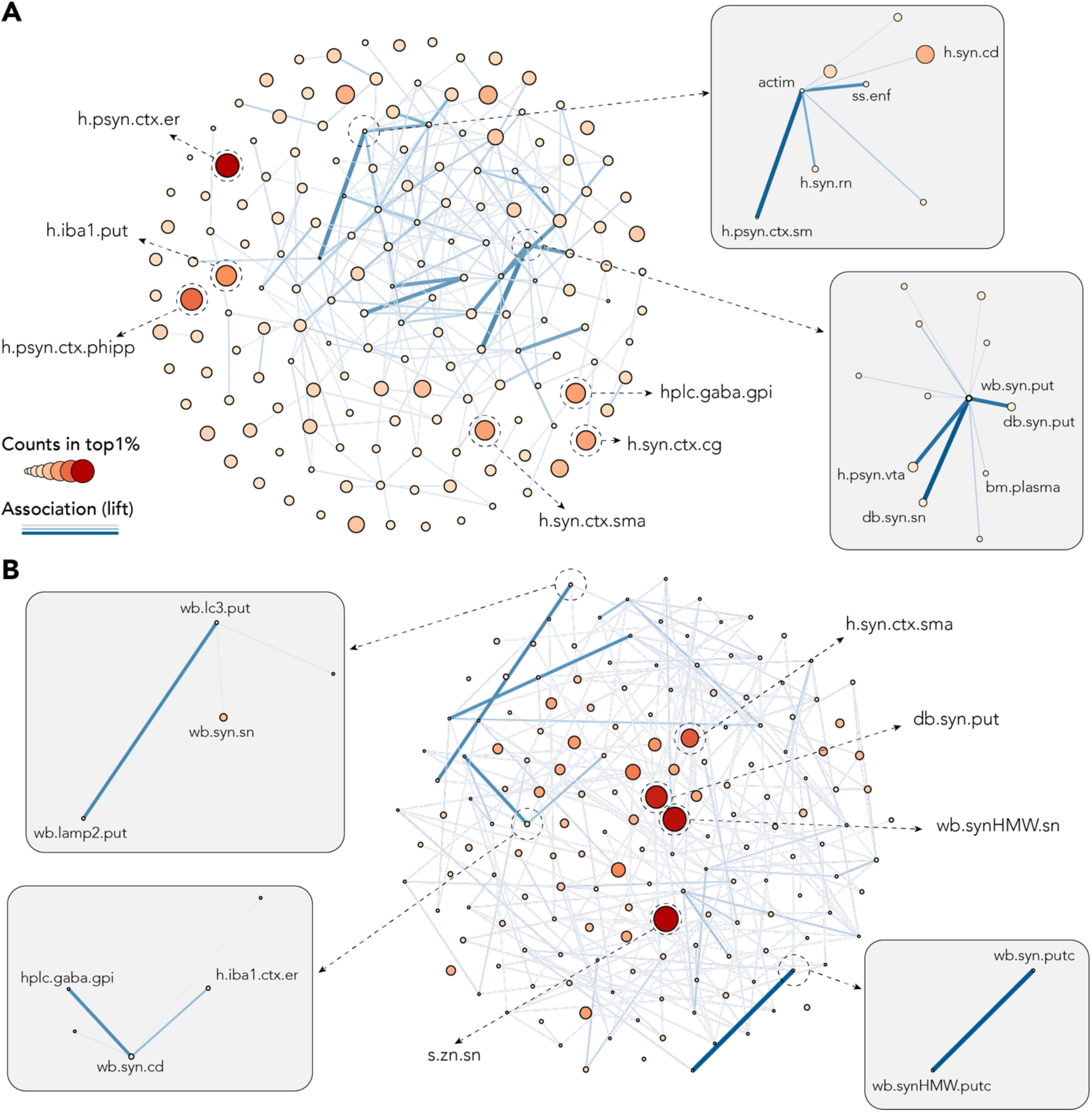
Association metric shows independence of strong predictors and beneficial association of weaker predictors. Both network plots were build using number of counts in the top1% as node size and color, and lift (association measure) as edges. To allow better visualization, only 10% of the strongest edges are shown. (**A**) Network plot for LB-injected animals showing independence of strong predictors: S129 phosphorylated α-syn (psyn) in the entorhinal (*h.psyn.ctx.er*) and the para-hippocampal cortex (*h.psyn.ctx.phipp*), microglia-activation in the putamen (*h.iba1.put*), α-syn in the cingulate cortex (h.syn.ctx.cg) and the supplementary motor area (*h.syn.ctx.sma*) and GABA levels in the internal part of the globus pallidus (*hlpc.gaba.gpi*). Upper right box highlights association between actimetry measure (*actim*) and a scan-sampling measure of body direction toward a closed environment (*ss.enf*) with α-syn levels in the caudate nucleus (*h.syn.cd*), the red nucleus (*h.syn.rn*) and psyn in the sensorimotor cortex (*h.psyn.ctx.sm*). Lower right box highlights association between pathological α-syn in the putamen (*wb.syn.put* and *db.syn.put*) and the SNpc (*db.syn.sn*) as well as psyn in the ventral tegmental area (*h.psyn.vta*) and peripheral levels of α-syn in the plasma (*bm.plasma*). (**B**) Network plot for noLB-injected animals showing independence of strong predictors: levels of Zn in the SNpc (*s.zn.sn*), pathological α-syn in the putamen (*db.syn.put*), α-syn in the supplementary motor area (*h.syn.ctx.sma*) and aggregated α-syn in the SNpc (*wb.synHMW.sn*). Upper left box highlights association between autophagosomes (*wb.lc3.put*) and lysosomes (*wb.lamp2.put*) levels in the putamen and α-syn in the SNpc (*wb.syn.sn*). Lower left box highlights association between GABA levels in the internal part of the globus pallidus (*hlpc.gaba.gpi*), α-syn in the caudate nucleus (*wb.syn.cd*) and microglia activation in the entorhinal cortex (*h.iba1.ctx.er*). Lower right box highlights association between soluble (*wb.syn.putc*) and aggregated (*wb.synHMW.putc*) levels of α-syn in the putamen.

In noLB-injected animals, the analysis shed light upon the relative abundance of two members of the macroautophagy pathway (Fig. 6B top left) as well as the balance between monomeric and high-molecular weight species of α-syn in the putamen (Fig. 6B bottom right). Such disruption of the nigrostriatal pathway has repercussions upon the basal ganglia physiology as GABA levels in their output structure, the internal globus pallidus, was associated with a decreased social behavior (Fig. 6B bottom left inset).

## Discussion

In the present study, we report that, in non-human primates, injection of distinct α-syn assemblies derived from PD patients lead to dopaminergic degeneration through discrete mechanisms. Applying a machine-learning method, we gained insight into unique signatures of degeneration induced by injection of two distinct α-syn pathogenic assemblies (i.e. those contained in the LB and noLB fractions derived from idiopathic PD patients’ brains). To do so, we built a large dataset with 180 variables obtained from behavioral, histological, biochemical, transcriptional and biophysical approaches applied to several brain areas for each individual. By using a distributed MLP algorithm that we developed for the purpose of this study, we identified characteristics that give insight into the strongest predictors of neurodegeneration for each experimental group. We have, therefore, described for the first time that distinct α-syn assemblies leading to similar degeneration in monkeys are associated with different mechanisms, hence experimentally confirming the true multifactorial nature of synucleinopathies.

Our results illustrate that both small oligomeric as well as larger α-syn assemblies induce dopaminergic degeneration in non-human primates. This finding was unexpected, since previous mouse studies from our laboratory showed that noLB injection did not have any observable consequence regarding dopaminergic degeneration, α-syn accumulation or phosphorylation (*11*). In agreement, other groups also showed the absence of toxicity of soluble recombinant α-syn (*12*).

One possible explanation is that primate dopaminergic neurons could be highly susceptible to α-syn toxicity. This could be in part due to their unique cellular architecture (*32*), a feature already known to contribute to the selective vulnerability of these neurons in PD (*33*). In fact, the large and complex axonal arbor of dopamine neurons make them particularly vulnerable to factors that contribute to cell death and, in primates, this axonal arbor is ten-fold the size of that in rodents (*32*). In addition, primate dopamine neurons display unique molecular characteristics (e.g. the presence of neuromelanin, the intracellular levels of which have been shown to be important in the threshold for the initiation of PD) (*34*). These unique features of primate dopaminergic neurons might be important in explaining the toxic mechanisms of the relatively low content of α-syn aggregates in the noLB fractions. Additional studies are now needed to fully address the question of host-seed interactions, but our results highlight the relevance and the need of the non-human primate model for the study of synucleinopathies.

We also confirmed that the toxicity mechanisms associated with patient-derived α-syn aggregates are shared features among patients and, therefore, common to the disease. Indeed, LB and noLB fractions used in this study were isolated from a pool of 5 patients who were different from the pool of 3 patients used in our previous study in mice (*11*). In the mice experiment (Fig. S3*B*) performed in this study, we observed the same level of dopaminergic degeneration (~40% at 4 months after injection).

The surprising observation, in non-human primates, that the noLB fraction is toxic to the same extent as the LB fraction suggests the existence of previously unrecognized forms of α-syn toxicity. Several studies have suggested that pre-fibrillar oligomeric species are the toxic α-syn species (*8*, *9*). Our biochemical studies showed that noLB and LB fractions had different amyloid properties (Fig. 1), contents (Fig. S1, S2A) and structures (Fig. S2B-E). Indeed, LB fractions contained a majority of large aggregated α-syn fibrils as well as some smaller aggregates while noLB fractions contained a smaller proportion (10 folds) of smaller aggregates and soluble α-syn. More importantly, the smaller aggregates were different in nature between LB and noLB fractions, as shown by micro-infrared spectroscopy (Fig. S2B-E). One could hypothesize that the observed effect is due to a species common between LB and noLB. However, because of the extent of degeneration, which was similar between the two experimental groups, and the α-syn content dissimilarity, both in amount and nature, this appears very unlikely. We believe that our results support the notion of the existence of a range of α-syn pathogenic structures with distinct toxic properties within the PD brain. Further work is necessary to provide a complete structural characterization of those species. As yet, very few studies report the high-resolution structures of α-syn aggregates, which are on the one hand, only derived from studies using recombinant α-syn and, on the other hand, limited to near atomic resolution (*35*–*37*). Encouragingly, much effort is currently being devoted to this field of research and two recent studies reported the atomic structure of α-syn fibrils determined by cryo-electron microscopy (*38*, *39*), while still being limited to recombinant-generated α-syn, and not isolated from human brain tissue.

In order to perform a characterization of the effects of the two fractions, we developed a machine learning method to identify their biological characteristics. It is now well accepted that machine learning algorithms can be trained to detect patterns as well as, or even better than, humans (*40*–*42*). Instead of the classification algorithms (the algorithm learns to identify in which category a sample belongs) that were mostly used in recent applications of machine learning in biology (*43*), we chose in this study to predict continuous and biologically-relevant variables using MLPs. Our choice was motivated by the limited sample size that is often a constraint of experimental biology. Although it might have been possible to use other feature selection methods, the use of MLPs with a distributed architecture allowed us to avoid overfitting issues and to develop a method particularly well-suited for low sample size datasets (*44*). As both LB and noLB-injected monkeys displayed similar levels of degeneration, they were indistinguishable using that endpoint. Instead of using a clustering analysis or a classification method, hence making the *a priori* assumption that these groups where different, we preferred to submit the two experimental groups to the MLP independently.

The combination of this constrained, distributed architecture and the holistic approach allowed us to rank input variables according to the number of times they appeared in the group of best predictors (defined as top 1% of best networks). A major issue in the use of machine learning in experimental biology in the ‘black-box’ is the fact that it is usually impossible to ‘understand’ how an algorithm predicted an output (*45*). By using a reverse engineering method, we aimed to tackle that issue. Because we explored all possible combinations of our variables, we could rank the input variables assuming that the more they appeared in the top 1%, the more they contained information allowing precise prediction of the neurodegeneration levels. Interestingly, our two experimental groups showed that some of the best predictors were similar (about 30%) but the majority were different. One could hypothesize that the similar variables between the two signatures probably embedded information that are consequences of neurodegeneration while the different ones probably contain information regarding the process of disease initiation and/or progression. Further experimental studies are now needed to confirm the relevance of these variables.

Also, as these two kinds of α-syn assemblies were associated with different signatures identified by our MLP approach, we propose that our results illustrate the multifactorial nature of the disease as different mechanisms (i.e. signatures) initiated by different triggers (i.e. α-syn assemblies) led to similar consequences (i.e. degeneration levels).

Using this methodology, we confirmed the interest of highly-expected variables but more importantly also unexpected variables that appear to be excellent predictors of α-syn-associated dopaminergic degeneration. The first hit for LB-injected animals was phosphorylated α-syn in the entorhinal cortex (as we have previously shown) followed by phosphorylated α-syn in the para-hippocampal cortex (unexpected), striatal microglial activation and GABA dysregulation in the internal part of the globus pallidus (expected) (Fig. S5). Conversely, Zn homeostasis was a strong predictive variable (unexpected) followed by α-syn aggregation-related terms (expected) in noLB-injected animals (Fig. S6).

In order to confirm the prediction made by the MLP approach, we first performed a retrospective literature analysis. This analysis showed that a significant part of the best predictors has been shown in the literature to be correlated with disease progression. Then, we attempted to confirm the interest of one of the top hits, the accumulation of Zn^2+^ in the SNpc, in independent experimental cohorts. Interestingly, we here describe that both in mice injected with noLB or in macaque monkeys (a different non-human primate sub species that the baboons used in that study) injected either in the striatum or in the SNpc, Zn levels were increased in the SNpc. However, in mice, Zn dyshomeostasis was not associated with neurodegeneration in the noLB group (at odds with what was observed in monkeys) suggesting a species difference in the relationship between zinc levels and dopaminergic tract integrity. Surprisingly, that result was not observed in rats and marmoset monkeys overexpressing human mutant α-syn. This observation might suggest that Zn dyshomeostasis is a feature of disease not triggered in the context of human mutant α-syn overexpression that is associated with fast progressing pathology (Bourdenx et al. 2015). Then, in order to expand our results to human pathology, we analyzed a publicly available proteomic dataset of human samples. According to that analysis, PD patients displayed increased levels of plasma membrane Zn transporters, hence suggesting a Zn dyshomeostasis in patients. In the context of PD, Zn dyshomeostasis has been associated with autophagy/lysosomal dysfunction in the context of *PARK9* mutations (Ramirez et al. 2006, Dehay et al. 2012). Further studies are now needed to fully unravel this connection.

Altogether, our findings show that primate dopaminergic neurons are sensitive to both small, mostly soluble, α-syn extracts as well as larger, aggregated, α-syn extracts derived from PD patients. These findings involve two immediate outcomes. First, since this toxicity has not been reported so far it suggest species differences that would need to be thoroughly investigated (*46*, *47*) and calls for a systematic appraisal of proteinopathies in primates in particular for validating therapeutic strategies before clinical testing (*48*). Second, the present study highlights the complex structure-toxicity relationship of α-syn assemblies and corroborates the multifactorial origin of synucleinopathies as distinct assemblies can induce similar degeneration (that would probably lead to similar clinical manifestation in patients) through different mechanisms, nigrostriatal or extranigral brain pathways, calling for molecular diagnosis to identify patient sub-populations before launching large-scale, heterogeneous in nature, clinical trials. Finally, we developed a machine-learning approach allowing and quantitative assessment of the explanatory power of a given set of variables compatible with the constrained sample size of experimental biology.

## Materials and Methods

### Access to data and machine-learning code for replicability and further use by the community

The entire raw data set is made available to the readers (Table S2). Authors chose not to provide representative examples of each procedure for the sake of space and because the entire data set is fully disclosed. Further information and requests for examples should be directed to and will be fulfilled by the Corresponding Contacts. Hyperlink to the machine-learning code (10.5281/zenodo.1240558) is provided (https://zenodo.org/record/1240558#.XC8pqy17Su4).

### Ethics statement

Experiments were performed in accordance with the European Union directive of September 22, 2010 (2010/63/EU) on the protection of animals used for scientific purposes. The Animal Experimentation Ethical Committee (CEEA) of the Vall d’Hebron Institute of Research (VHIR) approved experiments under the license number CEEA 81/13 (rats). The Institutional Animal Care and Ethical Committee of Bordeaux University (CE50, France) approved experiments under the license number 5012099-A (mice). The Institutional Animal Care and Ethical Committee of Murcia University (Spain) approved experiments under the license number REGA ES300305440012 (monkeys).

### Animals and Stereotactic Injections

#### Mice

Wild-type C57BL/6 mice (4 months old) received 2μl of either LB fractions or noLB fractions by stereotactic delivery to the region immediately above the right substantia nigra (coordinates from Bregma: AP=-2.9, L= −1,3, DV=-4.5) at a flow rate of 0.4μl/min and the pipette was left in place for 5 min after injection to avoid leakage. Mice were killed four months after injection. Ten to fifteen mice were used in each group.

#### Monkeys

Animals, whuch were from the research animal facility of the University of Murcia (Murcia, Spain) and housed in 2 multi-male multi-female exterior pens, were studied in a breeding farm over 2 years (Murcia, Spain). Animals were fed fruits, vegetables and monkey pellets twice a day before 9 am and after 5pm. Water was available ad libitum. 17 healthy adult olive baboons (P*apio papio*) were used in this study. Group sizes were chosen assuming a one-tailed alpha of 0.05, with sample size of at least three per group, which provided >80% power to detect a difference between the treatment groups and the control group, using a Fisher’s exact test. Animals were randomized into treatment or control groups. Six baboons were used for LB injections, four were used for noLB injections and seven were untreated control animals. Intrastriatal injections of either LB fractions or noLB fractions were performed at 2 rostrocaudal levels of the motor striatum (anterior commissure [AC], −1mm and −5mm) under stereotactic guidance as previously described (*49*–*52*). The total injected volume per hemisphere was 100μl (2 injection sites with 50μl each at 3μl/min at each location site). After each injection, the syringe was left in place for 10 min to prevent leakage along the needle track. A number of parameters were monitored during the course of the two-year study, including survival and clinical observations. At the end of the experiment (24 months post-injection), all monkeys were euthanised with pentobarbital overdose (150mg/kg i.v.), followed by perfusion with room-temperature 0.9% saline solution (containing 1% heparin) in accordance with accepted European Veterinary Medical Association guidelines. Brains were removed quickly after death. Each brain was then dissected along the midline and each hemisphere was divided into three parts. The left hemisphere was immediately frozen by immersion in isopentane at −50°C for at least 5 min and stored at −80°C. The right hemisphere was fixed for one week in 10 vol/tissue of 4% paraformaldehyde at 4°C, cryoprotected in two successive gradients of 20 then 30% sucrose in phosphate buffered saline (PBS) before being frozen by immersion in isopentane (−50°C) for at least 5 min and stored at −80°C until sectioning. CSF and blood samples (plasma, serum, whole blood) in the 17 animals were carefully collected before euthanasia. No samples were excluded from analysis in these studies.

### Purification of Lewy bodies from human PD Brains

The samples were obtained from brains collected in a Brain Donation Program of the Brain Bank “GIE NeuroCEB” run by a consortium of Patients Associations: ARSEP (association for research on multiple sclerosis), CSC (cerebellar ataxias), France Alzheimer and France Parkinson. The consents were signed by the patients themselves or their next of kin in their name, in accordance with the French Bioethical Laws. The Brain Bank GIE NeuroCEB (Bioresource Research Impact Factor number BB-0033-00011) has been declared at the Ministry of Higher Education and Research and has received approval to distribute samples (agreement AC-2013-1887). Human SNpc was dissected from fresh frozen postmortem midbrain samples from 5 patients with sporadic PD exhibiting conspicuous nigral LB pathology on neuropathological examination (mean age at death: 75 ± 2.75 years; frozen post-mortem interval: 31.8 ± 7.45h; GIE Neuro-CEB BB-0033-00011). Tissue was homogenized in 9 vol (w/v) ice-cold MSE buffer (10 mM MOPS/KOH, pH 7.4, 1Msucrose, 1mM EGTA, and 1mMEDTA) with protease inhibitor cocktail (Complete Mini; Boehringer Mannheim) with 12 strokes of a motor-driven glass/teflon dounce homogenizer. For LB purification, a sucrose step gradient was prepared by overlaying 2.2 M with 1.4 M and finally with 1.2 M sucrose in volume ratios of 3.5:8:8 (v/v). The homogenate was layered on the gradient and centrifuged at 160,000 x g for 3 h using a SW32.1 rotor (Beckman). Twenty-six fractions of 1500 μl were collected from each gradient from top (fraction 1) to bottom (fraction 26) and analyzed for the presence of α-synuclein aggregates by filter retardation assay, as previously described (*11*). Further characterization of LB fractions was performed by immunofluorescence, α-synuclein ELISA quantification and electron microscopy as previously described (*11*). For stereotactic injections, LB-containing fractions from PD patients were mixed together in the same proportion (PD#1, fractions 19 and 20; PD#2, fractions 19 and 20; PD#3, fraction 22; PD#4, fractions 17,18 and 19; PD#5, fractions 20, 21 and 23). NoLB-containing fractions (i.e. fraction 3, at the beginning of the 1,2M interface) derived from the same PD patients (which contain soluble or finely granular α-synuclein) but lacks large LB-linked α-synuclein aggregates were obtained from the same sucrose gradient purification. Using enzyme-linked immunosorbent assay (ELISA) kit against human α-synuclein (Invitrogen, #KHB0061 – following manufacturer’s recommendations), α-syn concentration was measured and both LB and noLB fractions were adjusted to ~24 pg α-synuclein per microliter. In all cases, samples were bath-sonicated for 5 min prior to *in vitro* and *in vivo* injections.

### Characterization of noLB and LB fractions

#### Electron microscopy

Briefly, carbon-coated nickel grids were covered for 1 min with corresponding fractions of interest, then washed 3 times with distilled water. They were then washed again in distilled water and stained for 5 min with 2% uranyl acetate, before being air-dried. Digital images were obtained with a computer linked directly to a CCD camera (Gatan) on a Hitachi H-7650 electron microscope. In all cases, samples were bath-sonicated for 5 min prior to the in vitro applications.

#### Immunofluorescence analysis of noLB and LB fractions

Indicated fractions from the sucrose gradient were spread over slides coated with poly-D lysine and fixed with 4% paraformaldehyde (PFA) in PBS for 30 min. Fixed slides were stained with 0.05% thioflavin S for 8 min and then washed three times with 80% EtOH for 5 min, followed by two washes in PBS for 5 min. Finally, all samples were washed 3 times with PBS and blocked with 2% casein and 2% normal goat serum for 30 min. For immunofluorescence analyses, samples were incubated with human α-synuclein specific antibody (clone syn211, Thermo Scientific, 1:1000) for 30 min, washed three times with PBS, incubated with a goat anti-mouse TRITC (Jackson, 1:500), before being cover-slipped for microscopic visualization using fluorescence mounting medium.

#### Dot-blotting analysis

To evaluate PK-resistant α-synuclein contained in noLB and LB fractions derived from PD brains, each fraction was subjected to digestion with 1 μg/ml proteinase K for 0, 15, 30, 45, and 60 min. The reaction was stopped by boiling for 5 min before dot-blotting with syn211 antibody. To analyze their stability, noLB and LB fractions were treated with increasing concentrations of urea (7 and 8M) or sodium dodecyl sulfate (SDS) (0.5, 1 and 2%) for 6 h at room temperature. α-Synuclein was visualized as described above.

Filter retardation assay of noLB and LB fractions were probed with antibodies against, phosphorylated α-synuclein (Abcam EP1536Y, 1:1000), ubiquitin (Sigma-Aldrich U5379, 1:1000), p62 (Progen GR62-C, 1:1000), hyperphosphorylated tau (AT8, MN1020, ThermoFischer) or A111111β (DAKO clone 6F/3D, 1:1000).

#### Human α -Synuclein aggregation TR-FRET immunoassay

Time-resolved Förster’s resonance energy transfer (TR-FRET)-based immunoassays were validated for total and oligomeric α-synuclein (*53*). Ten microliters of noLB and LB samples were analyzed for total α-synuclein quantification with the TR-FRET immunoassays kit against human α-synuclein aggregation kit (Cisbio, #6FASYPEG) according to the manufacturer’s instructions.

#### Velocity sedimentation and density floatation α-synuclein profiles in noLB and LB fractions

Frozen noLB and LB fractions aliquots (100 μL) were thawed and solubilized in solubilization buffer (SB) to reach 10 mM Tris pH 7.5, 150 mM NaCl, 0.5 mM EDTA, 1 mM DTT, Complete EDTA-free protease inhibitors (Roche), PhosSTOP phosphatase inhibitors (Roche), 1 U/μL Benzonase (Novagen), 2 mM MgCl_2_ and 2% (w/v) N-lauroyl-sarcosine (sarkosyl, Sigma) final concentrations, by incubating at 37°C under constant shaking at 600 rpm (Thermomixer, Eppendorf) for 45 minutes. For velocity sedimentations, a volume of 400 μL of solubilized noLB / LB fraction was loaded on top of a 11 mL continuous 5-20% iodixanol gradient (Optiprep, Sigma) in SB buffer containing 0.5% w/v final sarkosyl concentration, linearized directly in ultracentrifuge 11 mL tubes (Seton) with a Gradient Master (Biocomp). For density floatation gradients, a volume of 400 μL of solubilized noLB / LB fraction was mixed to reach 40% iodixanol in SB buffer with 0.5% w/v final sarkosyl concentration and loaded within an 11 mL 10-60% discontinuous iodixanol gradient in SB buffer with 0.5% w/v final sarkosyl concentration. The gradients were centrifuged at 180,000 g for 3 hours (velocity) or for 17 hours (density) in a swinging-bucket SW-40 Ti rotor using an Optima L-90K ultracentrifuge (Beckman Coulter). Gradients were then segregated into 16 equal fractions from the top using a piston fractionator (Biocomp) and a fraction collector (Gilson). Fractions were aliquoted for further analysis of their content by dot-blot. Gradient linearity was verified by refractometry.

For dot blotting, aliquots of the collected native fract α-synuclein immunolabelling, a step of fixation in PBS − 0.1% glutaraldehyde was performed at this point, followed by 3 washes in PBS. Membranes were then blocked with 5 % (w/v) skimmed milk powder in PBS − 0.1% (v/v) Tween and probed with anti-human α-synuclein (MJFR1, rabbit 1:10000, Abcam), anti-phospho pS129 α-synuclein (EP1536Y, rabbit 1:5000, Abcam) or anti α-synuclein aggregate specific FILA-1 (MJFR14-6-4-2, rabbit 1:10000, Abcam) primary antibodies in PBS-T − 4% (w/v) BSA, and secondary goat anti rabbit IgG HRP-conjugated antibodies (1:10000, Jackson Laboratories) in PBS-T 1% (w/v) milk. Immunoreactivity was visualized by chemiluminescence (GE Healthcare). The amount of the respective protein in each fraction was determined by the Image Studio Lite software, after acquisition of chemiluminescent signals with a Chemidoc imager (Biorad). Profiles obtained by immunoblot were normalized and plotted with SEM using the Prism software.

#### FTIR microspectroscopy

1-2 μL of each suspension was deposited on a CaF_2_ window and dried at room pressure and temperature. The protein aggregates were then measured in transmission at 50×50 μ^2^ spatial resolution with an infrared microscope (*54*). Depending on its size it was possible to collect one to twenty spectra inside each aggregate. The infrared microscope was a Thermo Scientific Continuum equipped with a MCT detector and a 32x 0.65 NA Reflachromat objective and matching condenser, coupled to a Thermo Scientific Nicolet 8700 spectrometer with a globar source and KBr beamsplitter. The microscope was operated in dual path single aperture mode. Spectra were recorded between 650-4000 cm^−1^ at 2 cm^−1^ resolution, with Happ-Genzel apodization and Mertz phase correction. Spectra were processed in Omnic 9.2 for automatic atmospheric correction to remove water vapor contribution.

### Rat Ventral Midbrain Primary Cultures

Postnatally derived ventral midbrain cultures were prepared essentially as previously described (*55*). Briefly, cultures were prepared in two steps. In the first step, rat astrocyte monolayers were generated as follows. The entire cerebral cortex from a single rat pup (postnatal days 1–2) was removed, diced, and then mechanically dissociated by gentle trituration. The isolated cells were plated at 80,000 cells per well under which a laminin-coated coverslip was affixed. The cells were housed at 37°C in an incubator in 5% CO_2_ and were fed on glial media (89% MEM, 9.9% calf serum, 0.33% glucose, 0.5 mM glutamine, and 5 μg/mL insulin). Once confluence had been attained (about 1 week in vitro), fluorodeoxyuridine (6.7 mg/mL) and uridine (16.5 mg/mL) were added to prevent additional proliferation. In the second stage, which occurred 1 week later, rat pups aged between 1 and 2 days were anesthetized and 1-mm^3^ blocks containing ventral midbrain neurons were dissected from 1-mm-thick sagittal sections taken along the midline of the brain. Tissues were collected immediately into cold phosphate buffer and were treated enzymatically using papain (20 U/mL) with kynurenate (500 μM) at 37°C under continuous oxygenation with gentle agitation for 2 h. A dissociated cell suspension was achieved by gentle trituration and was then plated onto the preestablished glia wells at a density of 0.5–1.7 million neurons per well. Cultures were maintained in specially designed neuronal media (47% MEM, 40% DMEM, 10% Hams F-12 nutrient medium, 1% calf serum, 0.25% albumin, 2 mg/mL glucose, 0.4 mM glutamine, 10 μg/mL catalase, 50 μM kynurenic acid, 10 μM CNQX, 25 μg/mL insulin, 100 μg/mL transferrin, 5 μg/mL superoxide dismutase, 2.4 μg/mL putrescine, 5.2 ng/mL Na_2_SeO_3_, 0.02 μg/mL triiodothyronine, 62.5 ng/mL progesterone, and 40 ng/mL cortisol) containing 27 μM fluorodeoxyuridine and 68 μM uridine to control glial outgrowth and in 10 ng/mL glial cell derived neurotrophic factor (GDNF). They were incubated for a further 7–8 days until the start of experiments. All tyrosine hydroxylase (TH) neurons were counted on each plate following the addition of noLB and LB fractions after 1, 2, 5 and 7 days of treatment.

### Non-Human Primate Behavioral Assessment

Following a 4-hour minimum habituation phase performed one day before the beginning of the observations, baboon behavior was observed outside the feeding and cleaning times, in a random order at two-time points (morning and afternoon), over 4 to 9 days (8 sessions per group). On the 1st observational time point (i.e. 1-month post-surgery), the habituation phase was performed over 3 days allowing the observer to recognize the animals individually. We used a scan-sampling method, appropriate for time budgeting (*56*), in which behavioral parameters were assessed every 5 minutes during 2-hour sessions, resulting in 192 scans per individual. Extra observational sessions were performed to avoid missing data. A unique trained observer (SC; intra-observer reliability: Spearman rank order correlation R=0.987) collected the data live on the 2-time points of the study: at 1- and 24-months post-surgery. The observer was standing 1 m away from the outdoor cages. We focused on behavioral profiles rather than single items and used two repertoires: one reports the interaction with the environment and one describes the position within the environment, according to published protocols (*57*–*59*). We investigated the percentages of occurrence of each item with regard to the total number of scans in order to obtain mean behavioral and postural time budgets, body orientation and location profiles.

### Histopathological analysis

#### Extent of lesion

To assess the integrity of the nigrostriatal pathway, tyrosine hydroxylase (TH) immunohistochemistry was performed on SNpc and striatal sections. Briefly, 50μm free-floating sections from one representative level of the striatum (anterior, medial and posterior) and serial sections (1/12) corresponding to the whole SNpc were incubated with a mouse monoclonal antibody raised against human TH (Millipore, MAB318, 1:5000) for one night at RT and revealed by an anti-mouse peroxidase EnVisionTM system (DAKO, K400311) followed by DAB visualization. Free-floating SNpc sections were mounted on gelatinized slides, counterstained with 0.1% cresyl violet solution, dehydrated and coverslipped, while striatal sections were mounted on gelatinized slides and coverslipped. The extent of the lesion in the striatum was quantified by optical density (OD). Sections were scanned in an Epson expression 10000XL high resolution scanner and images were used in ImageJ open source software to compare the grey level in each region of interest: i.e. caudate nucleus and putamen. TH-positive SNpc cells were counted by stereology blind with regard to the experimental condition using a Leica DM6000B motorized microscope coupled with the Mercator software (ExploraNova, France). The substantia nigra was delineated for each slide and probes for stereological counting were applied to the map obtained (size of probes was 100×80μm spaced by 600×400μm). Each TH-positive cell with its nucleus included in the probe was counted. The optical fractionator method was finally used to estimate the total number of TH-ositive cells in the SNpc of each monkey hemisphere. In addition, we measured Nissl cell count, the volume of SN, and the surface of TH-occupied in SN to fully characterize the pattern of dopaminergic cell loss in the SN.

#### α-synuclein pathology

Synucleinopathy was assessed with a mouse monoclonal antibody raised against human α-synuclein (syn211) and phosphorylated α-synuclein (clone11A5, Elan, 1:5000) immunostaining as we previously reported (*11*, *30*). Briefly, selected sections at two rostro-caudal levels were incubated in a same well to allow direct comparison of immunostaining intensity. Sections were incubated overnight at room temperature with the aforementioned antibodies. The following day, revelation was performed with anti-specie peroxidase EnVision system (DAKO) followed by 3,3’-diaminobenzidine (DAB) incubation. Sections were then mounted on gelatinized slides, dehydrated, counterstained if necessary and coverslipped until further analysis. Grey level quantification or immunostaining-positive surface quantification in forty brain regions (Fig. 2*B*) were performed as previously described (*30*).

#### Inflammation

Inflammatory process in the striatum, in the entorhinal cortex and in the white matter of noLB and LB-injected monkeys was measured through GFAP/S-100 (DAKO, Z0334/Abnova, PAP11341) and Iba1 (Abcam, ab5076) immunohistochemistry. Striatal sections of all animals were incubated together over night with a mix of rabbit antibodies raised against GFAP and S-100 for the astroglial staining (respective dilutions 1:2000 and 1:1000) and with a goat anti-Iba1 antibody for the microglial staining (dilution 1:1000). These signals were reveled with anti-specie peroxidase EnVision system (DAKO) followed by DAB incubation. Sections were mounted on slides, counter-stained in 0.1% cresyl violet solution, dehydrated and cover-slipped. Sections stained by GFAP-S-100 were numerized at x20 magnification with a NanoZoomer (Hamamatsu, France) and the quantification of GFAP-positive astrocytic reaction was estimated by a immunostaining-positive surface quantification at regional levels with the Mercator software (ExploraNova, France). Sections stained by Iba1 were used for the microglial morphology analysis through fractal dimension quantification based on microscopic acquisitions, as previously described(*60*). All analyses were performed blinded to the researcher.

### mRNA extraction and qRT-PCR

Substantia nigra samples were homogenized in Tri-reagent (Euromedex, France) and RNA was isolated using a standard chloroform/isopropanol protocol(*61*). RNA was processed and analyzed following an adaptation of published methods(*62*). cDNA was synthesized from 2 μg of total RNA using RevertAid Premium Reverse Transcriptase (Fermentas) and primed with oligo-dT primers (Fermentas) and random primers (Fermentas). QPCR was perfomed using a LightCycler® 480 Real-Time PCR System (Roche, Meylan, France). QPCR reactions were done in duplicate for each sample, using transcript-specific primers, cDNA (4 ng) and LightCycler 480 SYBR Green I Master (Roche) in a final volume of 10 μl. The PCR data were exported and analyzed in an informatics tool (Gene Expression Analysis Software Environment) developed at the NeuroCentre Magendie. For the determination of the reference gene, the Genorm method was used(*63*). Relative expression analysis was corrected for PCR efficiency and normalized against two reference genes. The proteasome subunit, beta type, 6 (Psmb6) and eukaryotic translation initiation factor 4a2 (EIF4A2) genes were used as reference genes. The relative level of expression was calculated using the comparative (2^−∆∆CT^) method(*63*).

Primers sequences: Psmb6 (NM_002798) forward: CAAGAAGGAGGGCAGGTGTACT; Psmb6 (NM_002798) reverse: CCTCCAATGGCAAAGGACTG; EIF4a2 (NM_001967) forward: TGACATGGACCAGAAGGAGAGA; EIF4a2 (NM_001967) reverse: TGATCAGAACACGACTTGACCCT; SNCA (CR457058) forward: GGGCAAGAATGAA GAAGGAGC; SNCA (CR457058) reverse: GCCTCATTGTCAGGATCCACA.

### Biochemical analysis

#### Total protein extraction and quantification

Immunoblot analyses were performed on substantia nigra, putamen and caudate nucleus. Five tissue patches were extracted on ice using 100μl of RIPA buffer (50 mM Tris-HCl pH 7.4, 150 mM NaCl, 1.0% Triton X-100, 0.5% Na-deoxycholate, 0.1% sodium dodecyl sulfate) with a protease inhibitor cocktail tablet (Complete Mini, Roche Diagnostics). The lysate was placed on ice for 20 min and then centrifuged at 14,000rpm for 15 min at 4°C. The supernatant was collected and the Bicinchoninic Acid (BCA) Assay was used to determine the total amount of protein in the lysates, and then stored at −80°C.

Based on total protein concentrations calculated from the BCA assays, aliquots of tissue lysates corresponding to known amounts of total protein per lane were prepared for each animal in Laemmli buffer (Tris-HCl 25mM pH=6.8, Glycerol 7.5%, SDS 1%, DTT 250mM and Bromophenol Blue 0.05%) for immunoblotting experiment.

#### Biochemical fractionation

This technique was performed as described(*64*). Tissue patches (n=10) were homogenized in 200μl of high-salt (HS) buffer (50 mmol/L of Tris, 750 mmol/L of NaCl, 5 mmol/L of EDTA, and a cocktail of protease inhibitors and phosphatase inhibitors). Samples were sedimented at 100,000 × g for 20 minutes, and supernatants were removed for analysis. Pellets were rehomogenized in successive buffers, after which each was sedimented, and supernatant was removed: HS containing 1% Triton X-100 (HS/Triton) (Variable names terminated as ultra.s1), RIPA (50 mmol/L of Tris, 150 mmol/L of NaCl, 5 mmol/L of EDTA, 1% NP40, 0.5% Na deoxycholate, and 0.1% SDS) (Variable names terminated as ultra.s12, and SDS/urea (8 mol/L of urea, 2% SDS, 10 mmol/L of Tris; pH 7.5) (Variable names terminated as ultra.p2). Sodium dodecyl sulfate sample buffer was added, and samples were heated to 100°C for 5 minutes prior to immunoblot analysis.

#### Western blot analysis

Western blots were run in all conditions from 20μg of protein separated by SDS-PAGE and transferred to nitrocellulose. Incubation of the primary antibodies was performed overnight at 4°C with rabbit anti-LC3 (1:1000, Novus Biologicals), rabbit anti-LAMP-2 (1:1000, Santa Cruz Biotechnology), mouse anti-TH (1:1000, Millipore), goat p62 (1:1000, Progen), mouse anti human-α-synuclein (1:1000, Thermo Scientific). For detection of ubiquitinated proteins, proteins were transferred on polyvinylidene fluoride membranes (Millipore) and subjected to Western blot analysis using a rabbit anti-Ubiquitin (1:1000, Sigma U5379). Anti-actin (1:5000, Sigma) was used to control equal loading. Appropriate secondary antibodies coupled to peroxidase were revealed using a Super Signal West Pico Chemiluminescent kit (Immobilon Western, Chemiluminescent HRP substrate, Millipore). Chemiluminescence images were acquired using the ChemiDoc+XRS system measurement (BioRad). Signals per lane were quantified using ImageJ and a ratio of signal on loading per animal was performed and used in statistical analyses.

#### Dot-blot analysis of α-synuclein

This technique was performed as we previously described(*9*, *11*). After heating at 100 °C for 5 min, 20 μg of protein extract was diluted in buffer (25 mM Tris-HCl, 200 mM Glycine, 1% SDS) and filtered through either a nitrocellulose membrane or an acetate cellulose membrane (Bio-Rad, 0.2 μm pore size). Membranes were then saturated in 5% dry-skimmed milk in PBS and probed with antibodies against α-synuclein (syn211, 1:1000), both α-synuclein fibrils and α-synuclein oligomers (Syn-O1, 1:10000(*65*, *66*)) (kindly provided by Prof. Omar El-Agnaf). Revelation was done as described in the previous Materials and Methods section.

### Synchrotron radiation X-ray fluorescence (SR-XRF) microscopy elemental mapping of brain tissue cryosections

The synchrotron experiments were carried out at Diamond Light Source, Harwell Science and Innovation Campus (Didcot, UK) with a 3 GeV energy of the storage ring and 300 mA currents with top-up injection mode. All SR-XRF microscopy investigations reported herein were carried out on the microfocus spectroscopy beamline (I18)(*67*). The micro X-ray fluorescence (μ-XRF) elemental mapping were acquired at room temperature with an incident X-ray energy set to 12 keV using an Si(111) monochromator and resulting in a X-ray photon flux of 2.10^11^ ph/s. The substantia nigra of each animal were collected from free-floating sections and mounted onto an X-ray transparent metal-free 4 μm thickness Ultralene ® foil (SPEXCert Prep, Metuchen, NJ, U.S.A.) secured to a customized Polyetheretherketone (PEEK) holder ensuring contamination-free samples and reduced X-ray scattering contribution. The samples were affixed to a magnetic plate that connects to the sample stage. The 4-element Si drift Vortex ME4 energy dispersive detector (Hitachi Hi-Technologies Science America) with Xspress-3 processing electronics, was operated in the 90° geometry, as such it minimizes the background signal. The sample-detector distance was fixed (75 mm). The sample was held at 45° to the incident X-ray beam and rastered in front of the beam whilst the X-ray fluorescence spectra were collected. An area of 500 μm x 500 μm within the substantia nigra pars compacta (SNpc) was mapped for each sample with a step-size that match the beam size (5 μm) and a dwell time of 1 s per pixel due to low concentration of the element. A thin (100 μm) pellet of the NIST standards reference materials SRM1577c (bovine liver material, NIST, Gaithersburg, MD, USA) was measured to calibrate experimental parameters as well as a thin-film XRF reference material (AXO Dresden GmbH). This was followed by elemental quantification through the open-source software PyMCA(*68*) in which both the reference material and the sample are modelled in terms of main composition, density and thickness. The fluorescence spectrum obtained from each pixel was fitted, the elemental concentration (μg/g dry weight or ppm) maps were generated and an average elemental concentration of the SNpc regions was obtained.

### Measurement of α-synuclein in monkey biological fluids samples

Multi-Array 96-well plates (MesoScale Discovery, Gaithersburg, MD, USA) were coated with 30μl 3μl/ml MJFR1 (abcam, Cambridge, UK) as capture antibody and incubated overnight at 4°C without shaking. The next day plates were washed 3 times with 150μl PBS-T [PBS (AppliChem, Darmstadt, Germany) supplemented with 0,05% Tween-20 (Roth, Karlsruhe, Germany)] per well. Unspecific binding of proteins was prevented by incubation with 150μl 1% BSA (SeraCare Life Sciences, Milford, MA, USA)/PBS-T/well for 1 hour and shaking at 700rpm. Calibrators (kindly provided by Prof. Omar El-Agnaf) were prepared from single use aliquots of α-synuclein (1μg/ml stored at −80°C until use) and ranged from 25000pg/ml to 6,1pg/ml in serial fourfold dilutions. 1% BSA/PBS-T served as blank. For the different specimen the following dilutions were applied: 1 in 10000 for whole blood and 1 in 8 for serum, plasma and CSF. All dilutions were prepared in 1% BSA/PBS-T. After washing the plates 25μl calibrator solutions and diluted samples were applied to the wells and incubated as indicated above. Plates were washed again and 25μl Sulfo-TAG labeled Syn1 antibody (BD Biosciences, Heidelberg, Germany) diluted to 1μg/ml in 1% PBS-T were applied to the wells as detection antibody. Sulfo-TAG labeling was done according to the manufacturer’s instruction using MSD Sulfo-TAG NHS-Ester (MSD). Incubation was for 1 hour at 700rpm. Plates were washed, 150μl 2x Read Buffer (MSD) was applied and the plates were read on a MSD SectorImager 2400. Data analysis was performed using WorkBench software (MSD).

### Neurotransmitter analysis

Brain patches were dissected out on ice-cold plate, weighed and put into 1.5 ml Eppendorf tubes. Samples were homogenized in methanol/water (50:50% v/v), then centrifuged at 14000 rpm for 15 min at 4°C(*69*). The supernatant was aliquoted and stored at −80°C until amino acid derivatization. Glutamate and GABA content in the samples was measured by HPLC coupled with fluorometric detection (FP-2020 Plus fluorimeter, Jasco, Tokyo, Japan) after precolumn derivatization with o-phthaldialdehyde/mercaptoethanol (OPA) reagent(*70*). Thirty microliters of OPA reagent were automatically added to 28 μL sample by a refrigerated autosampler kept at 4C° (Triathlon, Spark Holland, Emmen, The Netherlands). Fifty microliters of the mixture were injected onto a 5-C18 Hypersil ODS column (3 × 100 mm; Thermo-Fisher, USA) perfused at 0.48 mL/min (Jasco PU-2089 Plus Quaternary Pump; Jasco, Tokyo, Japan) with a mobile phase containing 0.1 M sodium acetate, 10% methanol, 2.2% tetrahydrofuran (pH 6.5). Chromatograms were acquired and analysed using a ChromNav software (Jasco, Tokyo, Japan). Under these conditions, the limits of detection for glutamate and GABA were ~1 nM and ~0.5 nM, and their retention times ~3.5 min and ~18.0 min, respectively.

### Multiple-Layer Perceptrons

Each Multiple-layer Perceptron (MLP) had the same architecture rule: 3 neurons as input, 3 neurons in the hidden layer and 3 neurons as output. Activation function of neurons was the hyperbolic tangent. Each network was trained over 1,000 presentations of a subset of the dataset. We used as error measure the mean square of differences between the expected output and the actual output. Our implementation comprises two parameters: a learning rate set at 0.05 (regulating the learning speed), and a momentum set at 0.05 (introducing purposefully a conservatism bias). Prior to learning, inputs were first scaled and centered (z scoring) in order to avoid dimensionality issues and then normalized between −0.5 and 0.5. For every combination of 3 variables used as inputs, 50 instances of MLP were trained with different subsets of the dataset. 80% of available data has been used for learning and the remaining 20% for testing the performance of the network (elements of each subset were randomly (and uniformly) drawn for each network). The performance from a given set of input variables was the mean of the error of the 50 instances of MLP that had data for these variables as inputs. Code was written using Python and the Python scientific stack(*71*–*73*) (Jones, 2001; Walt, 2011; Hunter, 2007). The code is fully available here (DOI: 10.5281/zenodo.1240558). Computation has been done using the Avakas cluster of the Mesocentre de Calcul Intensif Aquitain (MCIA). Rank-rank hypergeometric overlap (RRHO) test was performed as previously described(*74*) using RRHO package (1.14.0) in R(*75*) on variable list after ranking between experimental groups. Plotting was made using matplotlib in Python environment. The association metric was based on lift calculation. Let a and b be the two variables and n_x_ the number of combinations including variable x and n the total number of combinations considered in the analysis. Lift calculation was then:

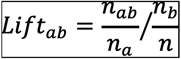

The lift calculation was then corrected for performance to avoid selection of detrimental association by being divided by the mean prediction error of the duo.

### Quantification and statistical analysis

Regarding the data analysis for FTIR microspectroscopy, spectra were analyzed by Principal Component Analysis (PCA). PCA is a multivariate statistical analysis technique that captures independent sources of variance in the data and represents them in Principal Components (eigenvectors) that carry the underlying spectral information and in a Score plot that shows the relation between spectra and can be used to cluster the data based on the spectral information. PCA were performed in The UnscramblerX 10.3 (Camo Software) using the SVD algorithm with leverage correction. Two series of preprocessing were applied prior to PCA and compared. Spectra were either baseline corrected in the amide I region between 1590 and 1700 cm^−1^ and vector normalized, or their second derivatives were computed and vector normalized.

Statistical analyses were performed with GraphPad Prism 6.0 (GraphPad Software, Inc., San Diego, CA). For all experiments, comparisons among means were performed by using One-way analysis of variance (ANOVA) followed, if appropriate, by a pairwise comparison between means by Tukey *post-hoc* analysis. All values are expressed as the meanm±standard error of the mean. Size effect was assessed with Cohen’s d analysis. In all analyses, statistical significance was set at p <0.05.

## Supporting information

Supplementary Informations

## Acknowledgments

The authors wish to express their gratitude to Pr. Alan R. Crossman (University of Manchester, UK) for his comments for his language supervision. We also thank Dr. Marion Bosc (Cold Spring Harbor, USA) for valuable comments on the manuscript. The authors thank Carmen Lagares Martínez (Head, Veterinary Service, University of Murcia) for administrative assistance; Maria Fermina Ros Romero and Josefa Martínez Rabadán (University of Murcia) for veterinary and husbandry support; Ana Luisa Gil, Lorena Cuenca and Ignacio Mascarell from Clinical and Experimental Neuroscience group (University of Murcia) for their technical help with various parts of the In Vivo part of these complex experiments. We would like to thank Dr. Philippe Hantraye (MIRCen) for providing baboon stereotactic frame. The University of Bordeaux and the Centre National de la Recherche Scientifique provided infrastructural support.

## Funding

This work was supported by a grant from the Michael J Fox Foundation (Project Grant No. 2013-8499), Fundacion de Investigacion HM Hospitales (Madrid, Spain), the Fundación Séneca (Project Grant No: FS19540/PI/14), the TARGET PD ANR grant and The Simone and Cino Del Duca Prize from French Academy of Sciences. MB and MLA were supported by a Ministère de l’Enseignement Supérieur et de la Recherche fellowship and the France Parkinson Foundation (MB). The help of the Bordeaux Imaging Center, part of the national infrastructure France BioImaging, granted by ANR-10INBS-04-0, is acknowledged. The Human α-Synuclein aggregation TR-FRET immunoassay was done in the Biochemistry and Biophysics Platform of the Bordeaux Neurocampus at the Bordeaux University funded by the LABEX BRAIN (ANR-10-LABX-43) with the help of Y. Rufin. Computing time for this study was provided by MCIA (Mesocentre de Calcul Intensif Aquitain), the public research HPC-center in Aquitaine, France. The samples were obtained from the Brain Bank GIE NeuroCEB (BRIF number 0033-00011), funded by the patients’ associations France Alzheimer, France Parkinson, ARSEP, and “Connaître les Syndromes Cérébelleux” to which we express our gratitude. The synchrotron Diamond is acknowledged for provision of beam time (exp. SP13009).

## Author contributions

M.B., M.V., J.O., P.D., B.D. and E.B. conceived and designed the study. M.B., G.P., I.T.D., C.E., N.G.C., M.T.H., B.D. and E.B. performed surgeries. S.C. and C.E. performed behavioral analysis. M.G. set up the actimetry behavioral platform. S.D., A.P. and P.A. performed histologic and immunohistochemical analysis of the data. S.D., A.P. and M.L.A. performed imaging experiments. E.D. performed electron microscopy analysis. F.L., M.L.A. and M.L.T. performed biochemistry experiments. C.P. performed and analyzed primary cultures experiment. S.B. and B.D. performed synchrotron analysis. C.S. performed infrared microscopy. N.K. and B.M. performed biological fluids analysis. S.N. and M.M. performed HPLC analysis. T.L.L. performed mRNA extraction and qPCR analysis. M.B., A.N., S.D., M.L.A., S.C., N.P.R., S.B., C.S., F.L., N.K., B.M., S.N., M.M., C.P., A.R., N.N.V. and O.E.A., M.T.H., P.D., M.V., J.O., B.D. and E.B. analyzed the data. M.B., A.N. and N.P.R. developed the MLP approach. M.B., M.V., J.O., B.D. and E.B. wrote the paper. B.D. and E.B. supervised the project. All authors discussed the results, assisted in the preparation and contributed to the manuscript. All authors approved the final version of the manuscript.

## Competing interests

E. Bezard is a director and a shareholder of Motac neuroscience Ltd. All the other authors have no conflict of interest to disclose.

## Supplementary Materials

**Table S1.**List of variables used in multiple-layer perceptron analyses.

**Table S2.**Raw data that served for the multiple-layer perceptron analyses for all behavioral, histological, biochemical, transcriptional and biophysical approaches (applied to several brain areas, totalizing the quantification of 180 variables for each individual).

## Notes

https://github.com/MathieuBo/Perceptromic

