## Supplementary Informations for "Identification of distinct pathological signatures induced by patient-derived α-synuclein structures in non-human primates"

**SUPPLEMENTAL INFORMATION**

**Supplemental Figures**

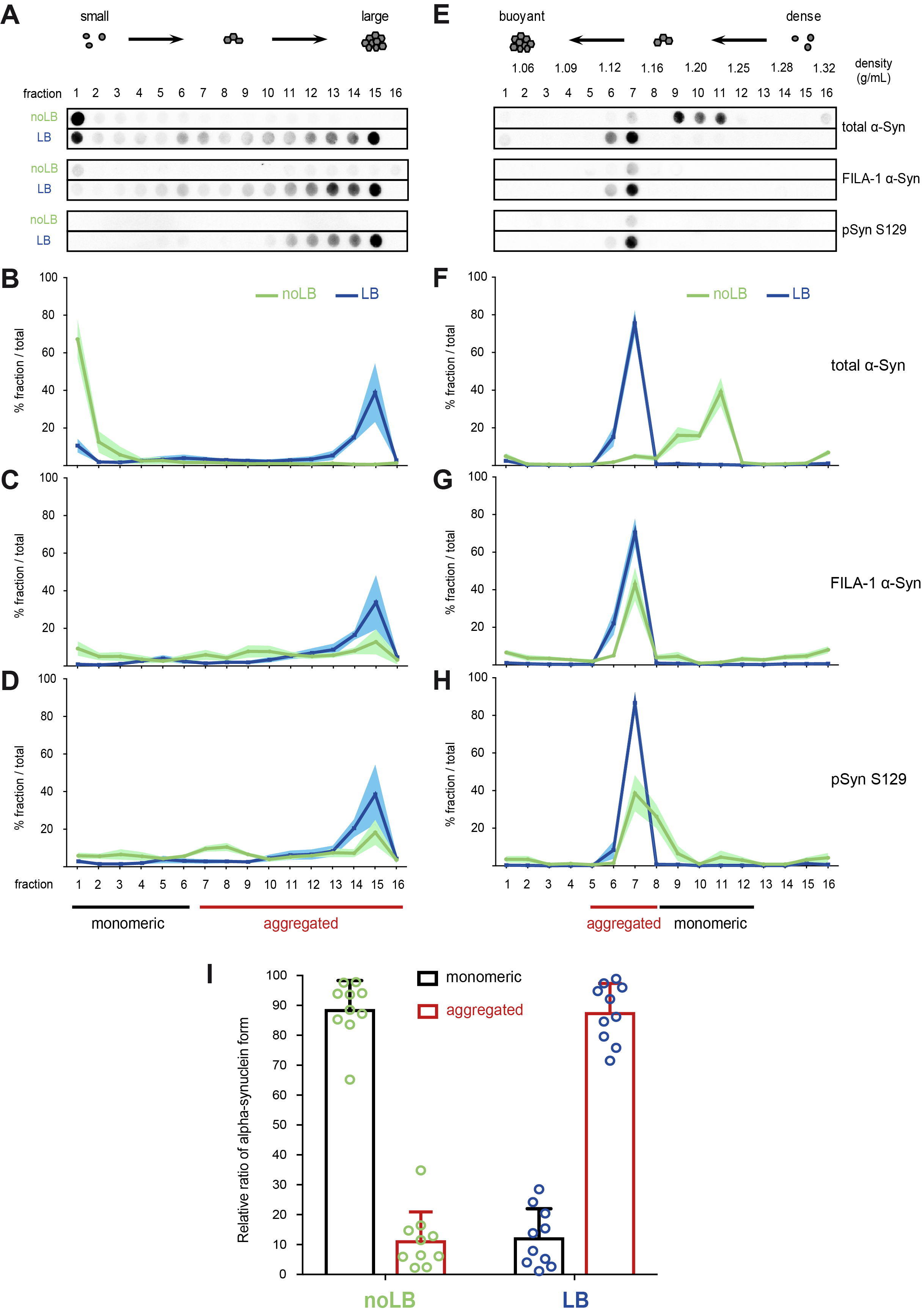

**Fig. S1. Relative quantification of soluble and aggregated** α-**synuclein in noLB/LB fractions by velocity sedimentation and density floatation gradient fractionations.** (**A**) Representative dot blots of the distribution of total α-synuclein (top, MJFR1 antibody, Abcam), FILA-1 α-synuclein aggregates (middle, MJFR14-6-4-2 antibody, Abcam) and phosphorylated pS129 α-synuclein (bottom, EP1536Y antibody, Abcam) on velocity sedimentation fractions (numbered from top to bottom of the gradient) for noLB (green) and LB (blue) fractions. (**B-D**) The relative amounts of total α-synuclein (**B**), FILA-1-positive α-synuclein (**C**) and pS129 α-synuclein (**D**) per fraction were quantified from noLB (green, n=5 PD patients) and LB (blue, n=5 same PD patients) velocity sedimentation fractionations. Mean curves (bold lines) with SEM (lighter shade areas) for each group. Velocity fractions containing soluble α-synuclein (1 to 6) are identified with a black line, while fractions containing aggregated insoluble α-synuclein (7 to 16) are identified with a red line. (**E**) Representative dot blots of total α-synuclein (top, MJFR1 antibody, Abcam), FILA-1 α-synuclein aggregates (middle, MJFR14-6-4-2 antibody, Abcam) and phosphorylated pS129 α-synuclein (bottom, EP1536Y antibody, Abcam) on density floatation gradient fractions (numbered from top to bottom of the gradient) for noLB (green) and LB (blue). (**F-H**) The relative amounts of total α-synuclein (**F**), FILA-1-positive α-synuclein (**G**) and pS129 α-synuclein (**H**) per fraction were quantified from noLB (green, n=5 PD patients) and LB (blue, n=5 same PD patients) equilibrium density floatation fractionations. Mean curves (bold lines) with SEM (lighter shade areas) for each group. Density fractions containing soluble α-synuclein (9 to 12) are identified with a black line, while fractions containing aggregated insoluble α-synuclein (5 to 8) are identified with a red line. (**I**) Relative ratios of soluble monomeric (velocity fractions 1-6 and density fractions 9-12) versus insoluble aggregated (velocity fractions 7-16, and density fractions 5-8) α-synuclein forms were calculated from each fractionation of noLB (green) and LB (blue) of each PD patient (n=5) and are represented respectively as black and red bars with SEM, together with individual values of each fractionation (n=5 velocities, n=5 densities).

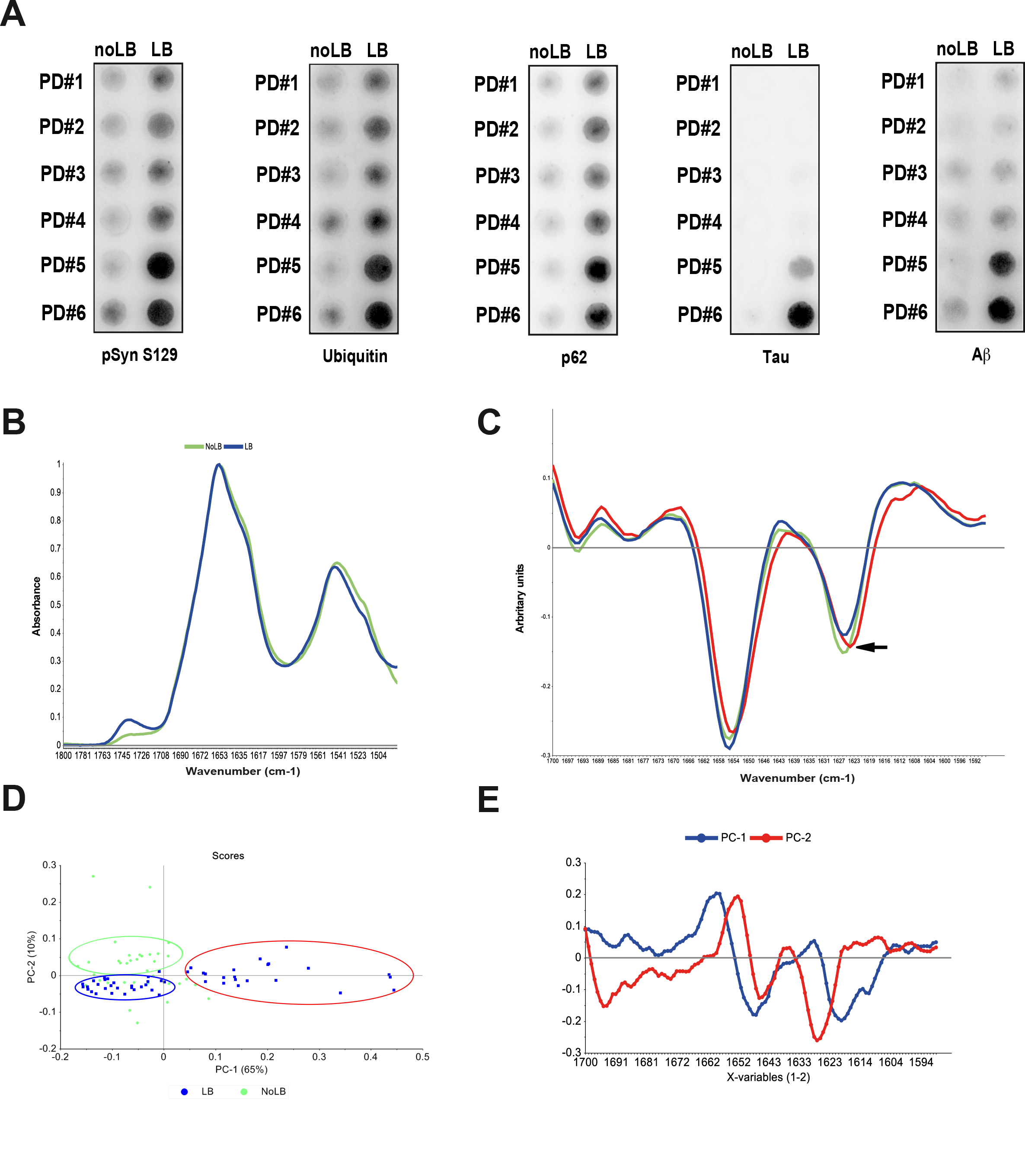
**Fig. S2. Analysis of the amyloid structure of LB and noLB protein aggregates by infrared microspectroscopy.** (**A**) Filter retardation assay of noLB and LB fractions probed with known components of LB: phosphorylated Ser129 α-syn (pSyn S129), ubiquitin, SQSTM1/p62, hyperphosphorylated tau and Aβ. Fractions were obtained by sucrose gradient fractionation from fresh frozen brain tissue from the sporadic PD patients used in this study (PD #1-5). (**B**). Mean spectra of LB (blue) and noLB (green) aggregates in the amide I and II bands. The spectra exhibited the typical amide I and amide II bands characteristic of protein samples. The amide I band in the two groups showed a strong shoulder at around 1630 cm^-1^ which is characteristic for the β-sheet component (**C**) Second derivative of the 3 groups separated by principal component analysis (PCA). A total of 37 no-LB and 53 LB vector normalized second derivative spectra were analyzed. The second derivative spectra allowed finding the exact position of the β-sheet component at 1626.7 cm^-1^. The 1627 cm^-1^/1653 cm^-1^ ratio does not completely separate the 2 groups by the intensity of their amyloid signal. One group containing about 40% of the LB spectra (red ellipse) presented a higher amyloid signal than the rest of the LB spectra and a 2 cm^-1^ shift in the position of the amyloid peak (black arrow) (**D**) PCA score plot showing the clustering of the spectra in 3 groups (in ellipses) formed by principal components 1 (PC-1) and 2 (PC-2). The PCA score plot shows that most LB and no LB spectra cluster on the negative part of PC-1 axis while over 40% of the LB spectra cluster on the positive part of PC-1 axis (red ellipse). In the negative part of PC-1 axis, LB and noLB groups can be separated along the PC-2 axis (in the green and blue ellipses). (**E**) PCA loadings of PC-1 and PC-2. Loadings of the PC-1 (blue) showing peaks at 1658, 1647 and 1620 cm^-1^ associated respectively with alpha-helices, random coil, and amyloid domains in the aggregates. Loadings of PC-2 (red) showing peaks at 1695, 1652, 1645 and 1628 cm^-1^ associated respectively with antiparallel beta-sheet, alpha-helix, random coil, parallel beta-sheet signal.

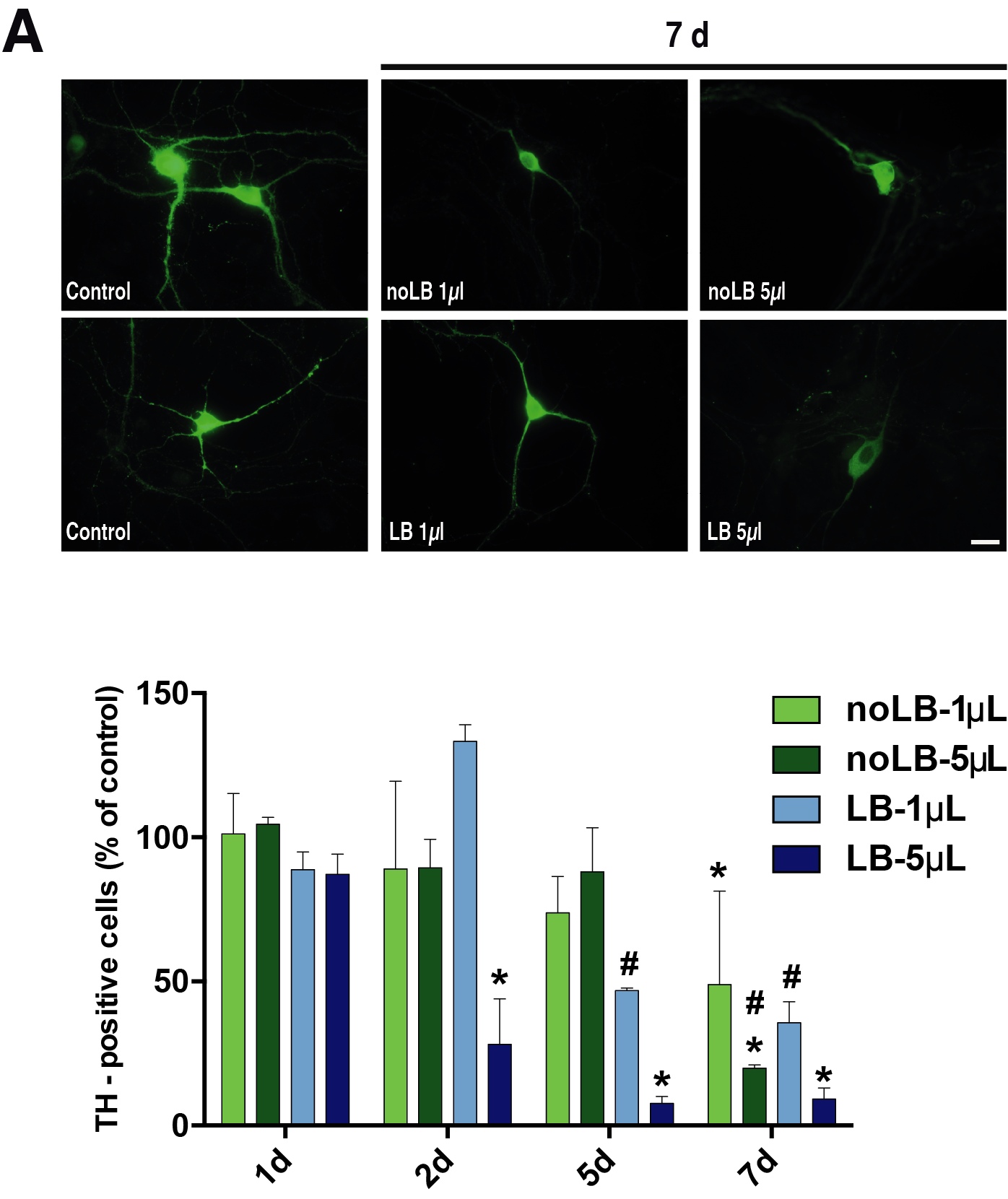

**Fig. S3. *In vitro* and toxicity of Lewy bodies (LB) and noLB inocula from Parkinson disease (PD) brains.** (**A, top**) In mouse primary mesencephalic culture, immunofluorescent labeling for tyrosine hydroxylase (TH) (green) following 7 days of treatment with 1µl and 5µl of noLB or LB fractions. Scale bars = 10µm. (**A, bottom**) Number of TH-positive primary mesencephalic neurons following the different treatments at 1, 2, 5 and 7 days. Analysis by Two-Way ANOVA followed by Tukey test for multiple comparisons. In all panels, n=2-6 per experimental group. *: p<0.05 compared with 1d; #: p<0.05 compared with 2d.

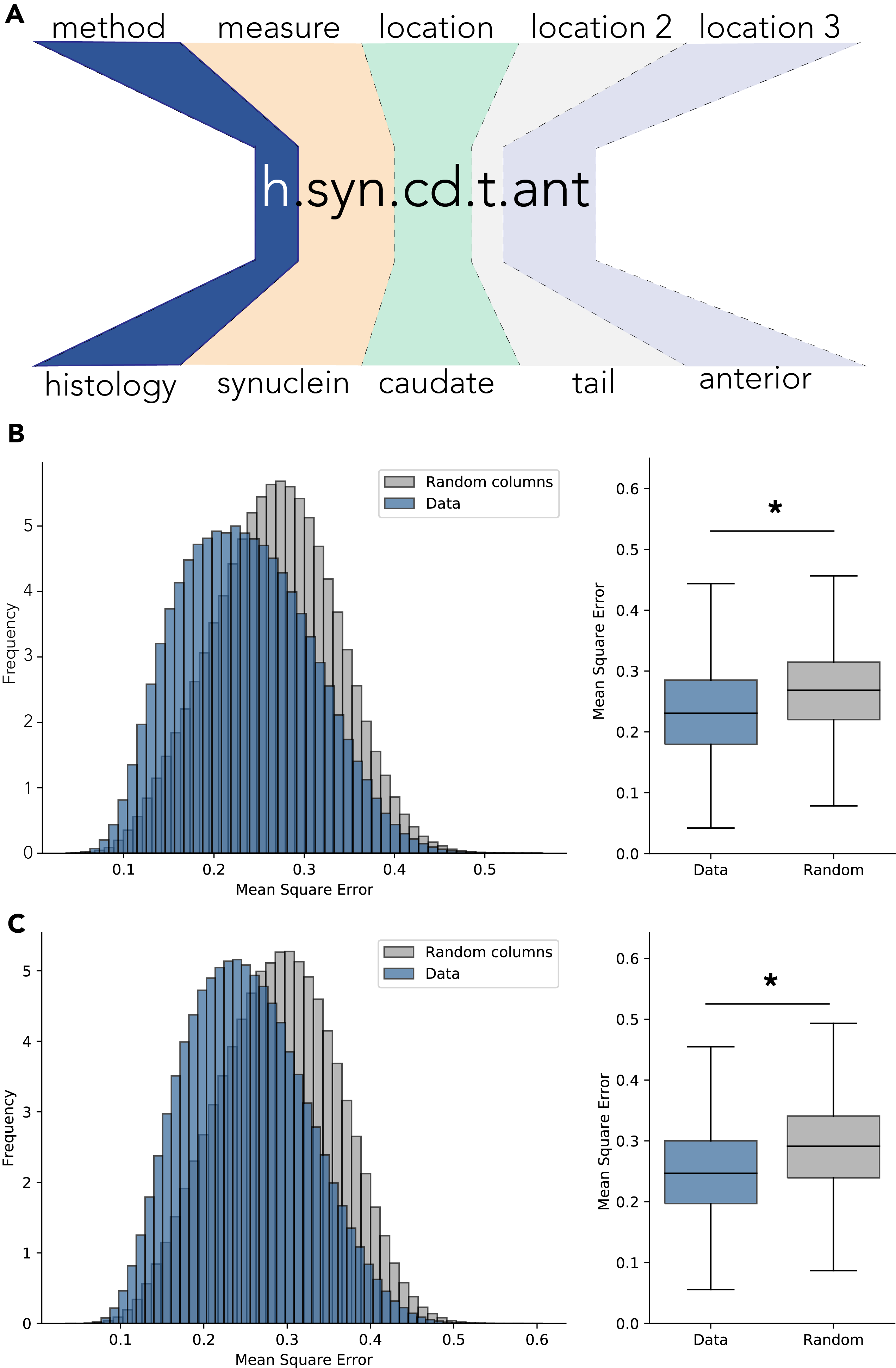

**Fig. S4. Variable nomenclature and performances of multiple-layer perceptrons. (A)** Guideline for variable naming. **(B, C)** A matrix of similar size filed with randomly generated values was used as control. (**B**) Data histogram for LB-injected group (left panel - blue) shows significant decrease (Cohen’s *d*= 0.468, t=-278.755, p<10^-6^) in mean square error. Right panel shows similar data presented as boxplots. (**C**) Data histogram for noLB-injected group (left panel - blue) shows significant decrease (Cohen’s *d*= 0.543, t=-323.013, p<10^-6^) in mean square error. Right panel shows similar data presented as boxplots.

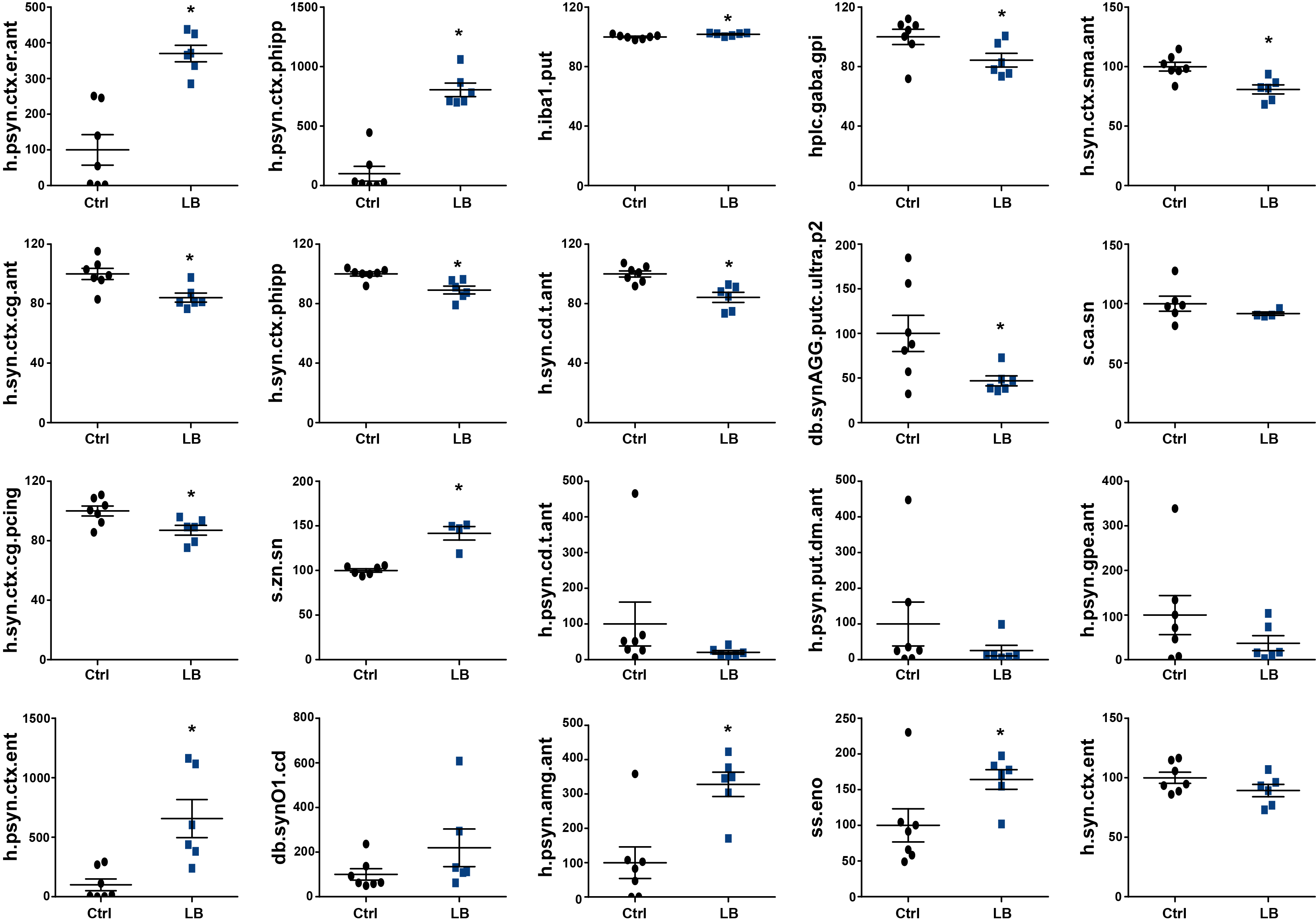

**Fig. S5. Pathological MLP-derived signature of LB-exposed monkeys.** Quantitative description of the 20 first-enriched variables for LB-injected animals highlighting a cortical-centric pattern: S129 phosphorylated α-syn (psyn) in the entorhinal (*h.psyn.ctx.er.ant*) [t _=_ 5.308; df = 11; p=0.0002] and the para-hippocampal cortex (*h.psyn.ctx.phipp*) [t _=_ 8.256; df = 11; p<0.0001], microglia activation in the putamen (*h.iba1.put*) [t _=_ 2.651; df = 11; p=0.023], GABA levels in the internal part of the globus pallidus (*hlpc.gaba.gpi*) [t _=_ 2.239; df = 11; p=0.047], α-syn in the supplementary motor area (*h.syn.ctx.sma.ant*) [t _=_ 3.574; df = 11; p=0.004], in the cingulate cortex (*h.syn.ctx.cg.ant*) [t _=_ 3.213; df = 11; p=0.008], in the para-hippocampal cortex (*h.syn.ctx.phipp*) [t _=_ 3.688; df = 11; p=0.004], and in the tail of the caudate nucleus (*h.syn.cd.t.ant*) [t _=_ 4.093; df = 11; p=0.002], pathological α-syn in the putamen (*db.synAGG.putc.ultra.p2*) [t _=_ 2.345; df = 11; p=0.038], levels of Ca in the SNpc (*s.ca.sn*) [t _=_ 1.058; df = 8; p=0.3211], α-syn in the posterior cingulate cortex (*h.syn.ctx.pcing*) [t _=_ 2.719; df = 11; p=0.020], levels of Zn in the SNpc (*s.zn.sn*) [t _=_ 6.389; df = 8; p=0.0002], S129 phosphorylated α-syn (psyn) in the tail of the caudate nucleus (*h.psyn.cd.t.ant*) [t _=_ 1.186; df = 11; p=0.261], S129 phosphorylated α-syn (psyn) in the dorsal part of the putamen (*h.psyn.put.dm.ant*) [t _=_ 1.091; df = 11; p=0.298], S129 phosphorylated α-syn (psyn) in the external part of the globus pallidus (*h.psyn.gpe.ant*) [t _=_ 1.261; df = 11; p=0.233] and in the entorhinal cortex (*h.psyn.ctx.ent*) [t _=_ 3.563; df = 11; p=0.005], aggregated α-syn in the caudate nucleus (*db.synO1.cd*) [t _=_ 1.452; df = 11; p=0.174], S129 phosphorylated α-syn (psyn) in the amygdala (*h.psyn.amg.ant*) [t _=_ 3.813; df = 11; p=0.003], a scan-sampling measure of body direction toward an open environment (*ss.eno*) [t _=_ 2.281; df = 11; p=0.043], α-syn in the entorhinal cortex (*h.syn.ctx.ent*) [t _=_ 1.550; df = 11; p=0.149]. The horizontal line indicates the average value per group ± SEM (n=7 from control animals; n=6 for LB-injected animals). Comparison were made using non-parametric t-test analysis. *p< 0.05.

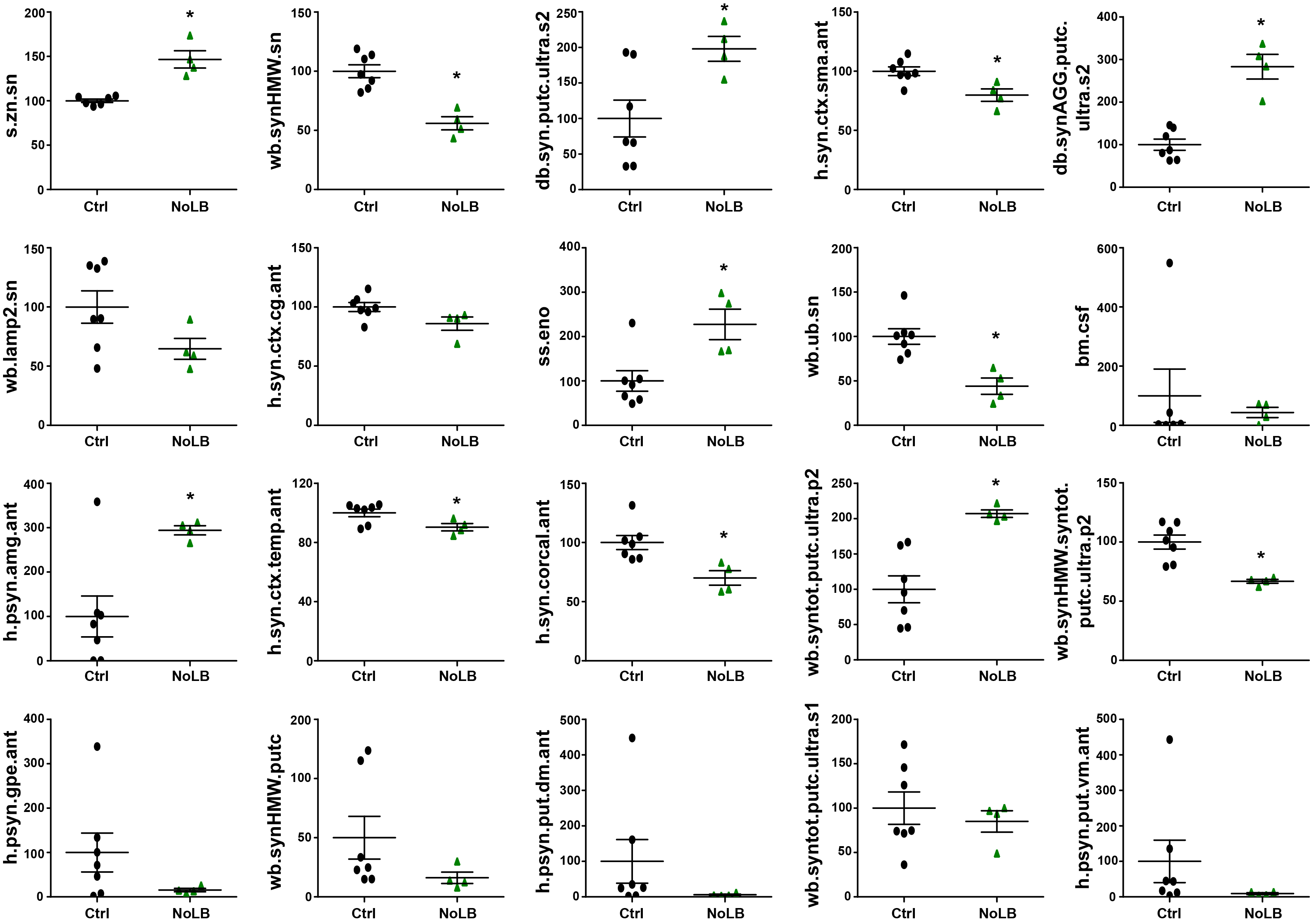

**Fig. S6. Pathological MLP-derived signature of noLB-exposed monkeys.** Quantitative description of the 20 first-enriched variables for noLB-injected animals illustrating a nigrostriatal-centric signature: levels of Zn in the SNpc (*s.zn.sn*) [t _=_ 5.735; df = 8; p=0.0004], aggregated α-syn in the SNpc (*wb.synHMW.sn*) [t _=_ 2.280; df = 9; p=0.043], pathological α-syn in the putamen (*db.syn.putc.ultra.s2*) [t _=_ 2.627; df = 9; p=0.028], α-syn in the supplementary motor area (*h.syn.ctx.sma.ant*) [t _=_ 3.194; df = 9; p=0.011], aggregated α-syn in the putamen (*db.synAGG.putc.ultra.s2*) [t _=_ 6.651; df = 9; p<0.0001], lysosomal levels in the SNpc (*wb.lamp2.sn*) [t _=_ 1.804; df = 9; p=0.105], α-syn in the cingulate cortex (*h.syn.ctx.cg.ant*) [t _=_ 2.148; df = 9; p=0.060], a scan-sampling measure of body direction toward an open environment (*ss.eno*) [t _=_ 3.182; df = 9; p=0.011], mono-ubiquitin levels in the SNpc (*wb.ub.sn*) [t _=_ 4.098; df = 9; p=0.003], α-syn levels in the CSF (*bm.csf*) [t _=_ 0.4991; df = 8; p=0.6311], S129 phosphorylated α-syn (psyn) in the amygdala (*h.psyn.amg.ant*) [t _=_ 3.081; df = 9; p=0.013], α-syn in the temporal cortex (*h.syn.ctx.temp.ant*) [t _=_ 2.451; df = 9; p=0.037] and in the corpus callosum (*h.syn.corcal.ant*) [t _=_ 3.249; df = 9; p=0.010], pathological α-syn in the putamen (*wb.syntotal.putc.ultra.p2*) [t _=_ 4.085; df = 9; p=0.003], aggregated α-syn in the putamen (*wb.synHMW.syntotal.putc.ultra.p2*) [t _=_ 4.085; df = 9; p=0.003], S129 phosphorylated α-syn (psyn) in the external part of the globus pallidus (*h.psyn.gpe.ant*) [t _=_ 1.428; df = 9; p=0.187], aggregated α-syn in the putamen (*wb.synHMW.putc*) [t _=_ 1.367; df = 9; p=0.205], S129 phosphorylated α-syn (psyn) in the dorsal part of the putamen (*h.psyn.put.dm.ant*) [t _=_ 1.131; df = 9; p=0.287], pathological α-syn in the putamen (*wb.syntotal.putc.ultra.s1*) [t _=_ 0.5697; df = 9; p=0.583], S129 phosphorylated α-syn (psyn) in the ventromedial part of the putamen (*h.psyn.put.vm.ant*) [t _=_ 1.124; df = 9; p=0.290]. The horizontal line indicates the average value per group ± SEM (n=7 from control animals; n=4 for noLB-injected animals). Comparison were made using non-parametric t-test analysis. *p< 0.05, when compared with control animals.

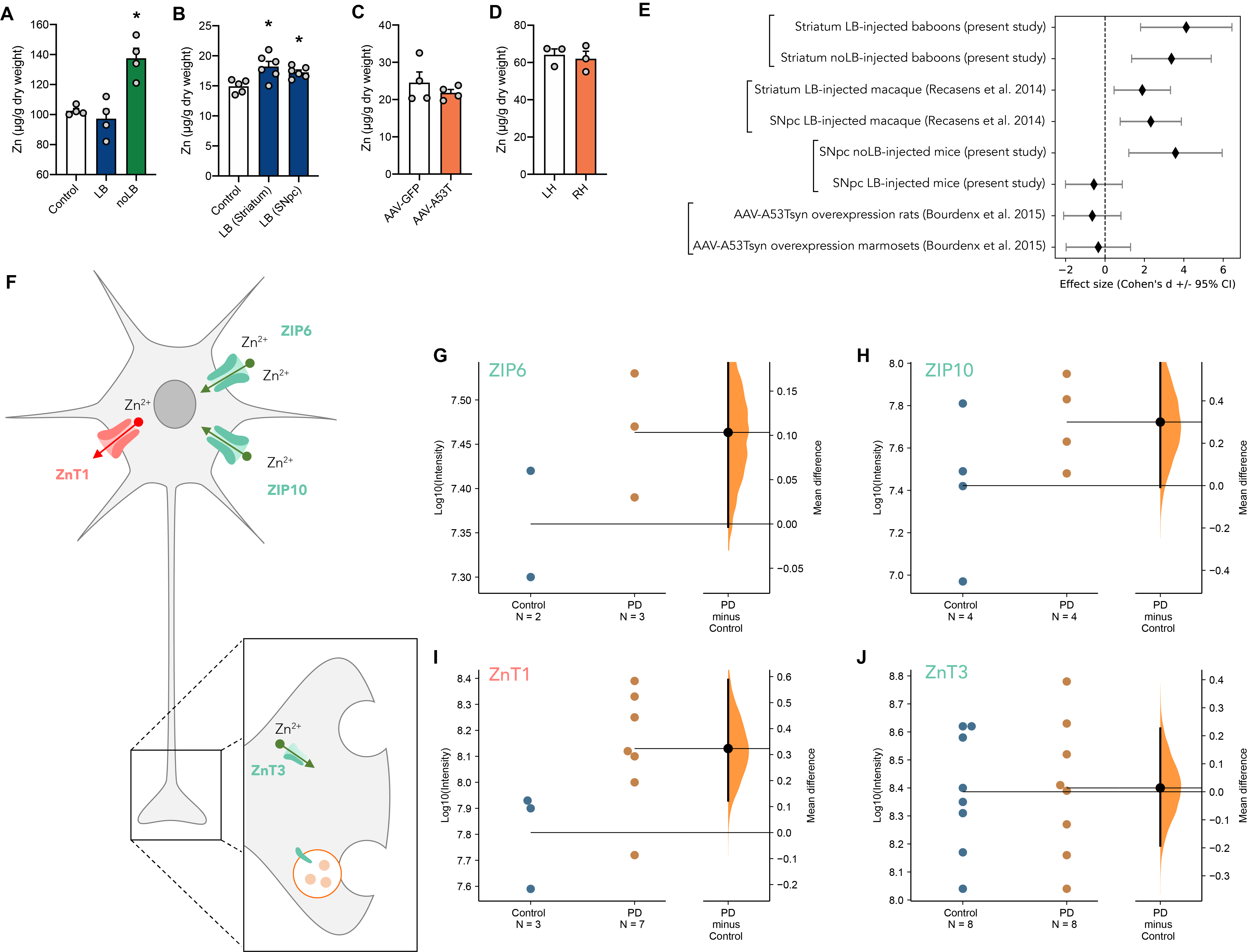

**Fig. S7. Nigral Zinc levels in independent experimental cohorts. (A)** Zinc levels in the substantia nigra of PD animal models in control, LB and noLB-injected mice (n=4 per group); (**B**) in control, striatum and SNpc LB-injected monkey macaques (5-6 nigral sections analyzed per animal); (**C**) in adeno-associated virus (AAV)-injected rats expressing either green fluorescent protein (AAV-GFP) or human mutant p.A53T α-synuclein (AAV-A53T) (n=4 per group); (**D**) in the left hemisphere (LH) or the right hemisphere (RH) of unilaterally injected marmoset monkeys (injection of AAV-A53T in the right hemisphere – n=3 per group). Data represent mean ± SEM. Comparisons were made using One-Way ANOVA and Sidak’s correction for multiple comparisons. *p< 0.05. (**E**) Metanalysis of the effect size of Zinc accumulation in the SNpc of animal cohort from the present study or from previous studies of our laboratory. Effect size is measured as the Cohen’s d +/- 95% confidence interval (CI). (**F**) Schematic drawing showing the localization and function of the different transporters identified in the study. Levels of (**G**) Zrt/Irt-like protein (ZIP) 6, (**H**) ZIP10, (**I**) Zinc transporter (ZnT) 1 and (**J**) ZnT3 in the frontal cortex of healthy individual versus PD/DLB patients (n=2-8 per group). Data are shown using Gardner-Altman plots. The right part of the plot indicates the mean difference +/- 95% confidence interval, the curve indicates the resampled distribution of mean difference.
